# Identification and characterization of an HtrA sheddase produced by *Coxiella burnetii*

**DOI:** 10.1101/2023.01.26.525556

**Authors:** Ikram Omar Osman, Aurelia Caputo, Lucile Pinault, Jean-Louis Mege, Anthony Levasseur, Christian A. Devaux

## Abstract

Having previously shown that soluble E-cadherin (sE-cad) is found in sera of Q fever patients, and that infection of BeWo cells by *C. burnetii* leads to modulation of the E-cad/β-cat pathway, our purpose was to identify which sheddase(s) might catalyze the cleavage of E-cad. Here, we searched for a direct mechanism of cleavage initiated by the bacterium itself, assuming the possible synthesis of a sheddase encoded in the genome of *C. burnetii* or an indirect mechanism based on the activation of a human sheddase. Using a straightforward bioinformatics approach to scan the complete genomes of four laboratory strains of *C. burnetii*, we demonstrate that *C. burnetii* encodes a 451 amino acid sheddase (CbHtrA) belonging to the HtrA family and differently expressed according to the bacterial virulence. An artificial CbHtrA gene (CoxbHtrA) was expressed and the CoxbHtrA recombinant protein was found to have sheddase activity. We also found evidence that the *C. burnetii* infection triggers an over-induction of the human HuHtrA gene expression. Finally, we demonstrate that cleavage of E-cad by CoxbHtrA on THP-1-cells leads to an M2 polarization of the target cells and the induction of their secretion of IL-10, which ‘disarms’ the target cells and improves *C. burnetii* replication. Taken together these results demonstrate that the genome of *C*. *burnetii* encodes a functional HtrA sheddase and establish a link between the HtrA sheddase-induced cleavage of E-cad, the M2 polarization of the target cells and their secretion of IL-10, and the intracellular replication of *C. burnetii*.

## Introduction

*C. burnetii* is an obligate intracellular pathogen that contaminates the host in the upper airways and is responsible for Q fever in humans(1). The bacteria are phagocytosed by pulmonary alveolar macrophages (myeloid cells) and *C. burnetii* replicates in late phagosomes(2). In non-myeloid cells, *C. burnetii* replication occurs in acidic phagolysosomal vacuole named CCV (for *C. burnetii-*containing vacuole), which require type IV secretion system (T4SS) to be fully established(3). In CCV the bacteria differentiate into the large cell variant and replicate, then the bacteria differentiate back into the infectious small cell variant and lyse the cell(4).Two to four weeks after bacterial exposure, the infected humans frequently present an acute fever (named Q fever). The symptoms of acute Q fever, usually resolves spontaneously in a few weeks, except in the 5% of patients who develop persistent Q fever(1, 5). Persistent Q fever is sometimes associated with endocarditis complications while interstitial lung diseases, persistent granulomatous hepatitis, cholecystis, and haemophagocytic syndrome have also been reported(6). Persistent Q fever patients have been considered to be at a higher risk of developing Non-Hodgkin lymphoma (NHL)(6–9). However, this last observation is controversial, and other studies of Q fever patients have not confirmed the link between Q fever and NHL(10, 11). If such a link did exist, it would be very rare and would result from a complex and long process of B-cell reprogramming.

Increasing evidence in the literature indicate that epithelial-cadherin (E-cad) can play a major role during the development of infectious diseases. During the human host invasion by bacteria, over-expression of proteases, called sheddases, can lead to intercellular adhesion molecule cleavage and tissue destruction leading to the transmigration of bacteria(12, 13). *Listeria monocytogenes* binds to intestinal epithelial cells through E-cad(14).Methylation of the *CDH1/E-cad* gene was frequently found in samples from *H. pylori* infected patients(15). *Chlamydia trachomatis* infection was also found to be associated with the methylation of the *CDH1* promoter and down-regulation of E-cad(16). High levels of sE-cad were reported in the biological fluids of patients with bacterial-associated tumors such as gastric adenocarcinoma induced by *Helicobacter pylori*(17), colon tumorigenesis associated with *Bacteroides fragilis* infection(18, 19), and colorectal tumors associated with *Streptococcus gallolyticus* infection(20). Changes in surface expression levels of adhesion molecules constitute a common mechanism for modulating cell-to-cell contact and cell reprogramming(21). Indeed, it was previously demonstrated that the tissue migration of immune-competent cells requires the sheddase-mediated degradation of transmembrane cell adhesion molecules (CAM)(22),whereas an over-expression of CAM can delay the rate of cell migration(23). Among cadherins, the E-cad, a 120 kDa cell-surface CAM, is known for its role in the cell-to-cell interaction crucial to ensuring the integrity of epithelia and cell polarity. Moreover, this trans-membrane protein behaves as a signaling molecule through its intra-cytoplasmic tail which binds second messengers (β-catenin), thereby playing a role in gene control, cell activation, division, differentiation and/or invasion(24). Cell reprogramming can be triggered by the release of soluble E-cadherin (sE-cad), an 80 kDa fragment generated from the proteolysis of the extracellular domain of E-cad(25). Elevated sE-cad concentrations in patients’ fluids were previously reported in association with a progression towards multiple myeloma and solid tumors(26).

We previously reported high levels of sE-cad in sera of patients infected by *C. burnetii*(27). We speculated that if there is a link between persistent Q fever and rare cases of NHL, one of the first steps towards NHL progression might be the over-expression of E-cad on monocytes, and the decrease of E-cad surface expression on the E-cad+CD20+ B-cells subpopulation of PBMCs (less than 1% of the B-cells) from *C. burnetii*-infected patients(27). In addition, because of the rapidly growing list of bacteria that mediate the proteolysis of E-cad to alter the integrity of epithelial tissues and initiate an invasion process(28), we also postulated that E-cad cleavage may be a step towards *C. burnetii-*invasion of the host during Q fever. Along with the discovery that sE-cad is released at high concentrations in the serum of *C. burnetii*-infected patients(27), the question arises about the nature of the molecule(s) catalyzing the cleavage of E-cad in those patients. Cleavage of the extracellular domain of E-cad has been reported to be achieved by a growing family of sheddases. Although elevated concentrations of the MMP-9 sheddase in sera of patients with acute Q fever and MMP-7 sheddase in sera of patients with persistent Q fever were previously reported(29, 30), it was tempting to postulate that prokaryotic sheddases encoded by *C. burnetii* could increase E-cad proteolysis(31), as previously reported for several bacteria(25, 32, 33) such as *Porphyromonas gingivalis*, *Bacillus subtilis*, *Bacteroides fragilis, Salmonella typhimurium*, and *Bacillus anthracis*.

We report here the results of an *in-silico* search for *C. burnetii* genes and proteins having homologies with known sheddases and provide the first direct evidence that *C. burnetii* can synthesize its own functional HtrA sheddase.

## Materials and methods

### Coxiella burnetii

Two laboratory strains of *Coxiella burnetii* were used in this study, the virulent Nine Mile I (NMI) and Guiana I (GuiI) and the isogenic Nine Mile and Guiana avirulent or “phase II” strain (NMII and GuiII) generated through serial *in vitro* passage, during which it undergoes a chromosomal deletion affecting the LPS biosynthesis. NMII and GuiII LPS lacks the unique branched terminal sugar-containing O-polysaccharide chain that is characteristic of the virulent strain (34, 35). *Coxiella burnetii* (Nine Mile strain RSA496, Guiana Strain Cb175) stocks were used to infect L929 cells. The L929 cells infected with either the Nine Mile or Guiana strain of *C. burnetii* cultured with a Minimum Eagle Medium (MEM, Invitrogen, USA) supplemented with 4% Fetal Bovine Serum (FBS, Invitrogen, USA) and 1% 2 mM L-Glutamine (Invitrogen, USA) were incubated at 35°C in a 5% CO_2_ atmosphere for the *in vitro* production of the bacteria as previously described(36). After two and five passages respectively for the Phase I and Phase II, the infected cells were sonicated, and the cell-free supernatants were centrifuged to harvest the bacteria, which were then washed and stored at −80°C until they were used. Gimenez staining and qPCR (using *com-1* gene) were used to estimate the concentration of bacteria in each sample.

### *In silico* sequence analysis

The genomes of four laboratory strains of *C. burnetii* were analyzed. The complete genomic sequences of RSA 493 Nine Mile(37), NL3262 Netherland(38), Z3055 clone(39) genotypically related to the strain causing the Netherlands outbreak, and Cb175 Guiana strain(9) were previously published and deposited in databases. The accession numbers for the genomic sequences of *C. burnetii* are: LK937696 *C* (Z3055 strain); NC_002971.4*C* (RSA 493 strain); NZ_CP013667.1 (3262 strain); and HG825990.3 (Cb175 strain). The protein-coding genes within these bacterial genomes were predicted using the open-source prodigal (PROkaryotic DYnamic programming Gene-finding ALgorithm) software tool(40). The different open reading frames encoding either known or hypothetical proteins of the 4 strains of *C. burnetii* were converted into protein sequences which were aligned at the protein level with known sheddases (22 sheddases, see below) in search of homologous sequences. Only genes showing significant e-values (<0.05) were kept for further investigation. Furthermore, we used the SignalP (Signal peptide) server for predicting signal peptides from amino acid sequences of proteins(41) to identify proteins compatible with secretion among the putative predicted sheddases from the 4 strains of *C. burnetii*.

Sequences of sheddases including zinc-dependent matrix metalloproteases, members of the disintegrin family, cysteine cathepsins, serine proteases, aspartic proteinases, bacterial proteases gingipains, subtilase serine protease, and high temperature requirement A protease(25) were obtained from the NCBI/NIH, GenBank and EMBL open international databases. Accession numbers for sequences of sheddases: NP_001289439.1 for gelatinase/MMP2; NP_002413.1 for stromelysine/MMP3; NP_002414.1 for matrilysin/MMP7; NP_004985.2 for gelatinase type IV collagenase/MMP9; NP_004986.1 for MMP14; NP_001101.1 for adamalysin/ADAM10; NP_997080.1 for ADAM15; NP_002644.4 for PITX1 pituitary homeobox; XP_002938616.2 for plasminogen; NP_001230055.1, NP_644806.1, and NP_005037.1 for the three isoforms of KLK7 kallikrein-7 isoform 3; AAA35655.1 for cathepsin; NP_689707.2 and NP_001137159.1 for the two isoforms of DCST1 E3 ubiquitin protein ligase; NP_872632.2, NP_872631.1, and NP_005218.1 for the three isoforms of EFNA4 ephrin-A4 isoform c precursor, AAB50410 for BFT *B. fragilis* toxin; 2LU1_A for Chain A Subtilase; ABJ97615.1 for rhomboid-1; and WP_005772880.1 for HtrA Do family serine endopeptidase. Once homologies were identified, sequences were further compared using Clustal Omega multiple sequence alignment (EMBL-EBI bioinformatic tool; Copyright EMBL 2020).

### Production of recombinant CbHtrA

A synthetic CoxbHtrA gene (length 1415 bp) cloned in the NdeI/NotI cloning site of the Novagen pET-22b(+) bacterial vector and optimized for *E. coli* expression, was purchased from GenScript Biotech (Leiden, Netherlands). The expressed protein was designed to include an N-terminal Strep-Tag WSHPQFEK (sequence of the tagged recombinant protein: MWSHPQFEKENLYFQSKKLSKIILSSIFAGLPLLLPVSSYAHLPSAVEGKTIPSLAPMLNKTTPSVVNIAVEKLIPQTPNPLQPEMDQNTAPTKVLGVGSGVIIDAKKGYIVTNAHVVKDQKIMVVTLKDGRRYRAKVIGKDEGFDLAVIQIHANHLTALPIGNSDQLKVGDFVVAVGSPFGLTQTVTSGVISALNRQEPRIDNFQSFIQTDAPINPGNSGGALIDLEGKLIGINTAIVTPSAGNIGIGFAIPSDMVKSVAEQLIKYGKVERGMLGVTAQNITPELADALNLKHNKGALVTKVVAESPAAKAGVEVQDIIESVNGIRIHSSAQLHNMLGLVRPGTKIELTVLRDHKVLPIKTEVADPKKVLLQRELPFLGGMRMQKFNDLEPDGTILQGVLVTGVDDSSDGALGGLEPGDIIISANGQLTPTVDELMKIAEGKPKELLLKVARGAGQLFLVIQQSQ of MW: 49,694 Da). Competent BL21(DE3) were transformed with the synthetic construct and grown in auto-inducing ZYP-5052 media at 37°C until it reached an O.D._600 nm_ of 0.6, then the temperature was set to 20°C for 20 hours. After centrifugation (5,000 g, 30 mins, 4°C), the resulting bacterial pellet was resuspended in 50 mM Tris pH 8, 300 mM NaCl and stored at − 80°C overnight. The bacterial extract was thawed and incubated on ice for 1 hour after adding lysozyme, DNAse I and PMSF (phenylmethylsulfonyl fluoride) to final concentrations of respectively 0.25 mg/mL, 10 μg/mL and 0.1 mM. The lysate was then submitted to sonication on a Q700 sonicator system (QSonica) and cells debris was discarded following a centrifugation step (12 000 g, 20 min, 4°C). The recombinant CoxbHtrA protein was purified using an ÄKTA avant system (GE Healthcare) equipped with a 5 mL StrepTrap HP column (GE Healthcare). The wash buffer contained 50 mM Tris pH 8, 300 mM NaCl and the elution buffer was 50 mM Tris pH 8, 300 mM NaCl, 2.5 mM desthiobiotin. Protein expression was assessed by SDS-PAGE and confirmed by performing MALDI-TOF MS analysis on gel bands previously obtained. The protein concentration was measured using a Nanodrop 2000c spectrophotometer (Thermo Scientific).

### *In vitro* cell culture model and infection

The human trophoblastic BeWo cell line was purchased from the American type culture collection (ATCC, CCL-98, Bethesda, USA) and cultured in a Dulbecco’s Modified Eagle Medium F-12 Nutrient Mixture (DMEM F-12, Invitrogen, USA) containing 10% Fetal Bovine Serum (FBS) (Invitrogen, USA). BeWo cells (2×10^5^ cells /well) were cultured in flat-bottom 24-well plates for 12 hours and were then infected with the live virulent and avirulent *C. burnetii* Nine Mile (NMI and NMII) strain and Guiana (GuiI and GuiII) strains, or were exposed to heat-inactivated virulent *C. burnetii* GuiI strains at a 50:1 bacterium-to-cell ratio, or stimulated with 100 ng/mL of lipopolysaccharide (LPS) from *E. Coli* (O55:B5; Sigma-Aldrich, USA) for 24 hours at 37°C in a 5% CO_2_ atmosphere.

THP-1 cells were cultured in RPMI-1640 containing 10% FBS, 2mM glutamine, 100U/mL penicillin and 50 μg/mL streptomycin and differentiated into macrophages after treatment with 50 ng/mL phorbol-12-myristate-13-acetate (Sigma Aldrich) for 48 hours(42, 43). Then THP-1 cells were polarized for 18 hours into the M1 phenotype in the presence of IFN-γ (20 ng/mL) and LPS (100 ng/mL), or into the M2 phenotype using IL-4 (20 ng/mL), or were maintained in the resting state without polarization (M0 phenotype)(44). The polarization status was confirmed by measuring the expression of M1 and M2 genes (**Table I**). As for infection experiments, THP-1 cells (2 × 10^5^ cells/well) were infected with the live virulent and avirulent *C. burnetii* Nine Mile strain (NMI and NMII).

**Table I:**
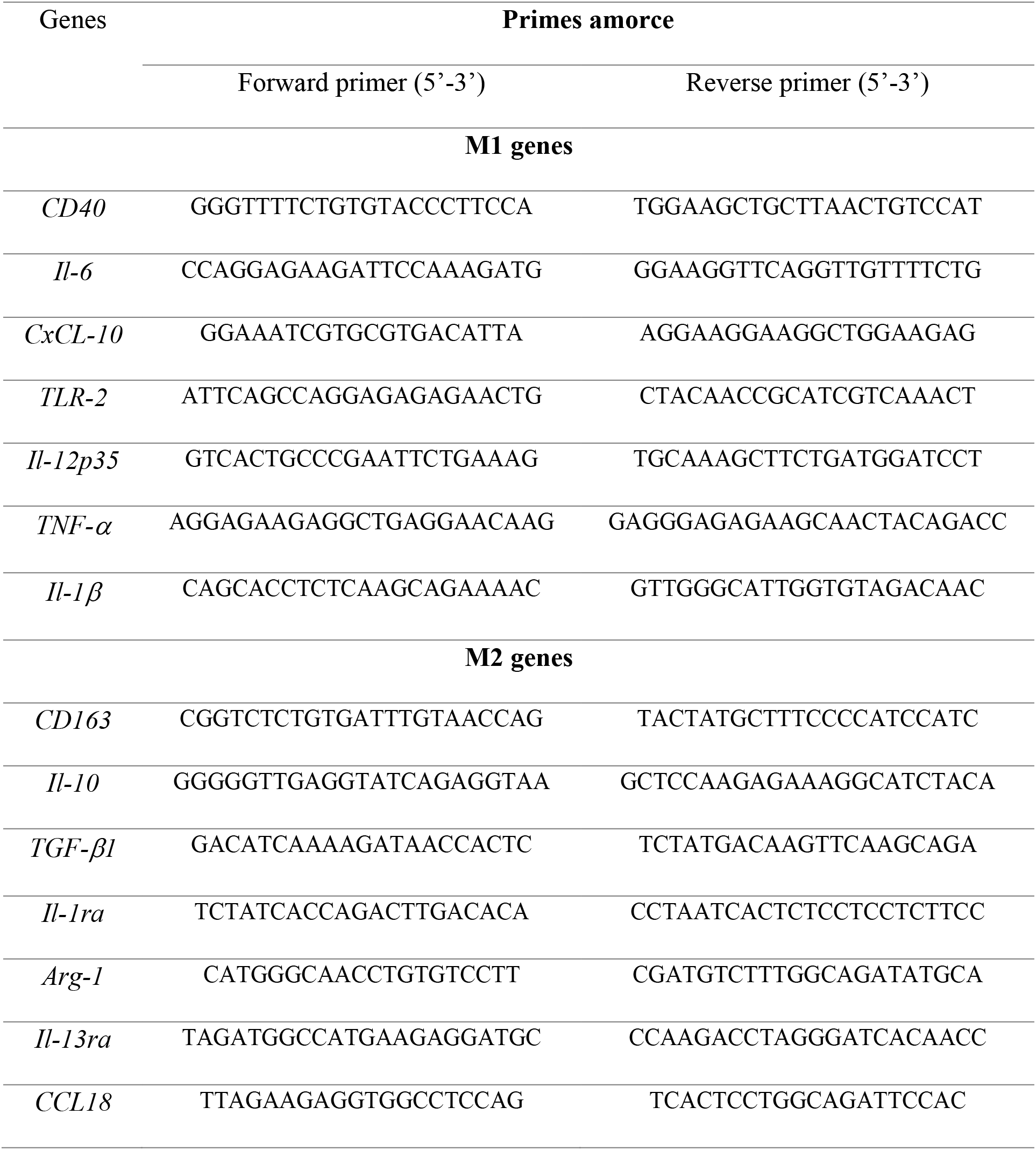
List of primers for inflammatory (M1)/ immunomodulatory (M2) genes

### DNA/RNA extraction and reverse transcription assay

Cellular and bacterial DNA were extracted using a Nucleic Acid purification kit (OMEGA, USA) according to the manufacturer’s instructions. The RNA of infected and uninfected cells was extracted using an RNeasy Mini Kit (QIAGEN SA, France) with a DNase I step according to the manufacturer’s instructions. Starting with 100 ng of purified RNA, the first-strand human cDNA was obtained using oligo(dT) primers and Moloney murine leukemia virus-reverse transcriptase (MMLV-RT kit; Life Technologies, USA), and The SuperScript™ VILO™ cDNA Synthesis Kit, for bacterial cDNA.

### Polymerase Chain Reaction (PCR) standard

The oligonucleotide primers designed (Fwd: 5’ GGCTCCCATTACTCCTGCC 3’; Rev: 5’ GAATCCGAATACCGTTGACAG 3’) for amplification of the CbHtrA gene allowed amplification of a DNA fragment of 897 bp corresponding to a portion of the gene in both strains. 5 μL of cellular/bacterial DNAs and cDNAs from infected and uninfected cells were used for the outer primer polymerase chain reaction. The 25 μL of polymerase chain reaction of each reaction included 12.5 μL of Master Mix AmpliTaq Gold™ 360 (Applied Biosystems, USA), 20 mM of each primer and 6 μL od DNAse/RNAse free water. The reaction was carried out in an Applied Biosystems MiniAmp Plus (Applied Biosystems, USA). The polymerase chain reaction was run at 95°C for 15 mins; (95°C for 30 seconds, 57°C for 30 seconds, 72°C for 52 seconds × 35 cycles) and 72°C for 5 mins.

### Quantitative Reverse Transcription-Polymerase Chain Reaction (qRT-PCR)

The qRT-PCR experiments were performed using specific oligonucleotide primers and hot-start polymerase (SYBR Green Fast Master Mix; Roche Diagnostics, Germany). The amplification cycles were performed using a C1000 Touch Thermal cycler (Biorad, USA). The specific primers used in this study were Human HtrA (*HuHtrA*) (Fwd: 5’ GAGATCACGTCTGGGAAGTC 3’; Rev: 5’ GAAAGTGACAGCTGGAATCTC 3’); *Coxiella burnetii* HtrA (*CbHtrA*) (Fwd: 5’ GGCTCCCATTACTCCTGCC 3’; Rev: 5’ CTAAGACTTTCGTTGGTGCTG 3’) and E-cadherin (*CDH1*) (Fwd: 5’ GAAGGTGACAGAGCCTCTGGAT 3’; Rev: 5’ GATCGGTTACCGTGATCAAAAT 3’). The results were normalized using the housekeeping gene β-actin (*ACTB*) (Fwd: 5’ CAT GCC ATC CTG CGT CTG GA 3’; Rev: 5’ CCG TGG CCA TCT CTT GCT CG 3’) and expressed as the relative expression (RE = 2^−ΔCT^) where ΔCt = Ct_Target_ − Ct_Actin_, and expressed as Fold Change (FC = 2^(−ΔΔCt)^) where ΔΔCt = [(Ct_Target_ − Ct_Actin_)_stimulated_] − [(Ct_Target_ − Ct_Actin_)_unstimulated_] as previously described(45). Ct values were defined as the number of cycles for which the fluorescence signals were detected.

### Evaluation of Coxb-HtrA functional activity

BeWo cells (2 × 10^5^ cells/well) were cultured in flat-bottom 12-well plates for 12 hours and were incubated with different dilutions (1:10, 1:100, 1:1000) of 1.6 mg/mL of CoxbHtrA recombinant protein. The recombinant protein elution buffer (1:10) (50 mM Tris pH 8, 300 mM NaCl) was used as a control. As well the cells were infected with the live virulent *C. burnetii* Nine Mile and Guiana strains or were exposed to heat-inactivated virulent *C. burnetii* strains at a 50:1 bacterium-to-cell ratio or stimulated with 100 ng/mL of lipopolysaccharide (LPS) from E. Coli (O55:B5; Sigma-Aldrich, USA). After 24 hours of infection and/or incubation in a 5% CO_2_ atmosphere, the culture supernatants were collected, centrifuged at 1,000 g for 10 mins and stored at −20°C until use. The quantity of sE-cad in the supernatants was determined using a specific immunoassay (DCADEO, R&D Systems, USA) according to the manufacturer’s instructions. The minimal detectable concentration of human sE-cad was 0.313 ng/mL. Protein quantification was evaluated using a western blot assay. For that purpose, cells were washed with ice cold phosphate buffered saline (PBS 1X) and lysed using the total protein extraction kit (NBP2-37853, Novus Biologicals, USA). Ten μg of protein was loaded onto 10% SDS polyacrylamide gels. After their transfer onto a Nitrocellulose membrane, the blots were incubated overnight at 4°C with a saturation solution (5% Free Fat Milk (FFM)-1X PBS - 0.3% Tween 20). Blots were then incubated (1:500 dilution) with mouse anti-human E-cad mAb (4A2C7, Life Technologies) and anti-human HtrA (PA523395, Life Technologies). The expression of Glyceradehyde-3-Phosphate dehydrogenase (GADPH) was measured using a mouse anti-human GADPH mAb (1:5000, Abnova, Taiwan) as the loading control. Afterwards the blots were incubated (1:5000 dilution) with a sheep anti-mouse horseradish peroxidase-conjugated antibody (Life Technologies, France). The proteins were revealed using an ECL Western Blotting Substrate (Promega, USA).

### Flow cytometry assay

BeWo cells (1×10^6^ cells/well) were cultured in flat-bottom 6-well plates and were then incubated in a 5% CO_2_ atmosphere for 24 hours with 1.6 μg/mL of the CoxbHtrA recombinant protein. Cells were exposed to 2 mM EDTA in PBS for 15 mins at 4°C to release the adherent cells from the culture plate. After centrifugation at 500 g for 5 mins, the pellet was suspended in a FACS buffer (2 mM EDTA, 10% FBS in PBS) and was then incubated (1:1000 dilution) for 1 hour with a mouse anti-human E-Cad ectodomain antibody (HECD-131700, 1:1000 dilution; Life Technologies). After two washes, cells were incubated with a goat anti-mouse IgG, Alexa Fluor-488 secondary antibody. Fluorescence intensity was measured using a Canto II cytofluorometer (Becton Dickinson/Biosciences, France) and the results were analyzed using a FlowJo V10.7.2 software (Becton Dickinson, USA).

### Confocal microscopy analysis

THP-1 macrophage cells were cultured on sterile coverslips in 24-well plates. After fixation with paraformaldehyde (3%), the cells were permeabilized with 0.1% Triton X-100 for three minutes and saturated with 3% BSA-0.1% Tween 20-PBS for 30 minutes at room temperature. Cells were incubated for one hour at room temperature with a mouse monoclonal anti-E-cadherin (4A2C7, Life Technologies, France) directed against the cytoplasmic domain of E-cad and revealed using an anti-mouse IgG (H+L) secondary antibody (Alexa Fluor 555) (Life Technologies). The 4’,6’-diamino-2-fenil-indol (DAPI) (1:2500, Life Technologies) and the Phalloidin (Alexa-488) (1:500, Ozyme) were used to stain the nucleus and the filamentous actin, respectively.

### Antimicrobial activity of macrophage-differentiated THP-1

THP-1-macrophages (2×10^5^ cells/well) were pretreated with CoxbHtrA (1μg/mL) for 24 hours before being infected with C. burnetii bacterium (bacterium-to-cell ratio of 50:1) for 4 hours. After extensive washing to remove free bacteria (time designed as T0), infected cells were cultured for 9 additional days. DNA (50μL volume) was extracted from the total infected cells/assay every 3 days using DNA Mini Kit (Omega, USA). Infection was quantified using 2 μL of DNA and real time quantitative PCR (qPCR) performed with specific primers targeting the *C. burnetii* com-1 gene(46). Bacterial uptake and their intracellular fate were expressed as the number of bacterial DNA copies within the THP-1-macrophage.

### Immunoassays

The release of IL-10 and IL-6 were quantified in cell supernatants using specific immunoassay kits purchased from BD Biosciences. The sensitivity of assays was 15.6 pg/mL and 4.68 pg/mL, respectively.

### Statistical analysis

The statistical analyses of the data were performed using the GraphPad-Prism software (version 9.0). The results are presented as the ± standard error of the mean (SEM). The Mann-Whitney U test was used for group comparison. A p-value<0.05 was considered statistically significant.

## Results

### *In silico* search for a hypothetical sheddase in the genome of *C. burnetii*

The proteolytic cleavage of the E-cad and release of sE-cad observed in BeWo cells after *C. burnetii* infection(31) suggests that the bacterial infection is accompanied by expression of sheddase(s). Since sheddases may be either of cellular origin or encoded by infectious pathogens, we first focused our attention to investigate a possible bacterial origin of the sheddase involved in releasing the sE-cad.

A bioinformatic strategy was set up in order to search for genes potentially coding sheddases in the genome of *C. burnetii*. First, the genes coding for 22 sheddases were blasted against the full genome (1,481 genes screened) from 4 strains of *C. burnetii* (RSA 493 Nine Mile; NL3262 Netherland, Z3055 clone related to the NL3262 strain and Cb175 Guiana strain), in search of homologous sequences. Because this early screening was unsuccessful, a similar investigation was conducted at the protein level using the sequences from: MMP2, -3, -7, -9, and -14; ADAM10, and -15; PITX1 pituitary homeobox; plasminogen; the three isoforms of KLK-7; cathepsin; the two isoforms of DCST1 E3 ubiquitin protein ligase; the three isoforms of EFNA4, BFT, *B. fragilis* toxin; and HtrA, *Salmonella typhimurium* high temperature requirement protein A. These protein sequences of sheddases were aligned against the different open reading frames (hypothetical gene products) of *C. burnetii*. Of 22 sheddases tested, only 3 human sheddases (HuMMP-9; HuADAM-15; HuMMP-3), 1 parasite sheddase (rhomboid), and 2 sheddases of bacterial origin (subtilase and HtrA) revealed significant alignment with hypothetical gene products from *C. burnetii*. Except for the bacterial HtrA, the other protein sequences matched with only a short portion of a known or hypothetical protein from *C. burnetii* (**Table II**).

**Table II:** Identification of proteins from *C. burnetii* matching with sheddase sequences using BLAST analysis. Only the conserved sequences presenting a significant e-value (e-value <0.05) are recorded in this table. The % identity means the percent of identical and related amino acids with respect to the sequence of the sheddase used for the BLAST. The % coverage refers to the percent identity found with respect to the length of the *C. burnetii* sequence studied. Only three *C. burnetii* hypothetical molecules found to share similarities with rhomboid, subtilase and HtrA carry secretion signal peptides (SIP).

| *C. burnetii* | Protein | Sheddases | % | E-value | % | Signal |
| --- | --- | --- | --- | --- | --- | --- |
| strain |  |  | Identity |  | Coverage | IP |
| **Cb175** | HG825990.3_1452 | MMP-3 | 28 | 5.10<sup>-5</sup> | 95 | No |
|  | HG825990.3_2153 | ADAM_15 | 30 | 7.10<sup>-4</sup> | 53 | No |
|  | HG825990.3_823 | subtilase | 31 | 3.10<sup>-5</sup> | 48 | Yes |
|  | HG825990.3_51 | Rhomboid-1 | 26 | 6.10<sup>-4</sup> | 61 | Yes |
|  | HG825990 3_847 | HtrA | 42 | 2e-112 | 100 | Yes |
| **NL3262** | NZ_CP013667.1_1221 | MMP-3 | 30 | 5.10<sup>-4</sup> | 45 | No |
|  | NZ_CP013667.1_2102 | MMP-9 | 25 | 0.001 | 79 | No |
|  | NZ_CP013667.1_1792 | Subtilase | 31 | 2.10<sup>-5</sup> | 48 | Yes |
|  | NZ_CP013667.1_230 | Rhomboid-1 | 26 | 6.10<sup>-4</sup> | 61 | Yes |
|  | NZ_CP013667.1_1768 | HtrA | 42 | 2e-112 | 100 | Yes |
| **RSA 493** | NC_002971.4_1204 | MMP-3 | 30 | 5.10<sup>-4</sup> | 45 | No |
|  | NC_002971.4_271 | MMP-9 | 25 | 0.001 | 79 | No |
|  | NC_002971.4_682 | Subtilase | 31 | 3.10<sup>-5</sup> | 48 | Yes |
|  | NC_002971.4_45 | Rhomboid-1 | 26 | 6.10<sup>-4</sup> | 61 | Yes |
|  | NC_002971.4_703 | HtrA | 42 | 2e-112 | 100 | Yes |
| **Z3055** | LK937696.1_1129 | MMP-3 | 30 | 5.10<sup>-4</sup> | 45 | No |
|  | LK937696.1_272 | MMP-9 | 25 | 0.001 | 79 | No |
|  | LK937696.1_695 | Subtilase | 31 | 2.10<sup>-5</sup> | 48 | Yes |
|  | LK937696.1_44 | Rhomboid-1 | 26 | 6.10<sup>-4</sup> | 61 | Yes |
|  | LK937696.1_716 | HtrA | 42 | 2e-112 | 100 | Yes |

Regarding the similarities with sheddases of human (Hu) origin, 3 strains of *C. burnetii* (RSA 493 Nine Mile, NL3262 Netherland, and Z3055) expressed a 79 amino acid sequence presenting 25% identity and 79% coverage with HuMMP-9 (expected (e)-value 10^−3^). This sequence matched with amino acids 539 to 660 of HuMMP-9 (a protein composed of 707 amino acid residues), a region corresponding to a hemopexin domain. The Guiana strain Cb175 (known to lack genes compared to the other strains) lacks expression of this *C. burnetii* (Cb) MMP-9-like protein sequence. In contrast, the Cb175 strain expresses a protein presenting 30% identity and 50% coverage with HuADAM-15 (e-value 7.10^−4^) that is not found within the genome of the other *C. burnetii* strains studied. Finally, all strains of *C. burnetii* encode a protein sequence showing 28-30% identity with the 1/3 NH-2 terminal region of the HuMMP-3 metalloprotease (a protein composed of 477 amino acid residues). Regarding the parasite and bacterial sheddases, sequence identity was found for rhomboid, subtilase and HtrA.

The blasting search in the NCBI database revealed that the CbMMP-3-like sequence corresponds to the nuclear transport factor 2 protein and that the CbMMP-9-like sequence corresponds to the L28 protein of the large 50S ribosomal subunit of *C. burnetii*. (**Table III**). Because the CbMMP-3 like/nuclear transport factor 2 protein shows unexpected similarities with HuMMP-3 in the region that corresponds to the catalytic domain of HuMMP-3 and shares some sequence similarities with the BFT toxin bacterial sheddase, we performed a reverse BLAST that lead us to the conclusion that it is very unlikely that such a molecule might exhibit sheddase activity. Using the same approach, we also eliminated CbMMP-9 like/ L28 protein and CbADAM-15 like as possible candidates for sheddase activity. Two additional hypothetical *C. burnetii* proteins with either Rif1 or OmpA domains emerged from the *in-silico* screening. However, none of those molecules met the criteria for a possible sheddase.

**Table III:** Identification of the sheddase-like proteins encoded by the *C. burnetii* genome. Blasting the sequences for homologies with the human proteins of MMP and ADAM families against sequences from the NCBI database revealed that the hypothetical CbMMP-3-like sequence corresponded to a nuclear transport factor 2 (NTF2) protein and shared some similarities with the BFT toxin bacterial sheddase, while the hypothetical CbMMP-9-like sequence corresponded to the L28 protein of the large 50S ribosomal subunit of *C. burnetii*. To investigate the relevance of these results, a reverse BLAST analysis was conducted in which the best matches between CbMMP-3 like/NTF2, CbMMP-9like/L28 protein and CbADAM15-like sequences from *C. burnetii* and the full human genome were investigated to identify the sequences producing the most significant alignments (not shown). The best matches for CbMMP-3 like/NTF2 were Homo sapiens cadherin 13 (e value of 0.39; percent identity of 87%; percent coverage of 7%), and Homo sapiens RAS p21 protein activator 2 (e value of 0.39; percent identity of 80%; percent coverage of 8%). The best matches for CbMMP-9 like/L28 protein were Homo sapiens Rho GTPase activating protein 20 (e value of 0.76; percent identity of 83%; percent coverage of 17%) and Homo sapiens synthrophin gamma 2 (e value of 0.76; percent identity of 92%; percent coverage of 11%). The best match for CbADAM-15 like protein was Homo sapiens cell division cycle 25 B (e value of 0.11; percent identity of 93%; percent coverage of 6%). Since no human gene found by the reverse BLAST analysis turned out to be a sheddase family member, it is very unlikely that these proteins from *C. burnetii* show sheddase activity. Using the same strategy we found that the CbRhomboid-1 corresponds to a protein with an Omp-A domain, and CbSubtilase-like corresponds to a protein with a Rif1 domain. Again, no human genes found by reverse BLAST analysis turned out to be a member of the sheddase. The only hypothetical protein found using our screening strategy which met the required criteria to possibly be a sheddase was CbHtrA

| **C. burnetii strains** | **Sheddases** | **Mean E-value** | **Mean % Coverage** | **Identification** |
| --- | --- | --- | --- | --- |
| **Cb175** | ADAM_15 | 7.10<sup>-4</sup> | 53 | Hypothetical *C. burnetii* protein |
| **NL3262; RSA 493; Z3055** | MMP-9 | 0.001 | 79 | 50s ribosomal protein L28 |
| **Cb175**<br>**NL3262**<br>**RSA 493**<br>**Z3055** | MMP-3 | 5.10<sup>-4</sup> to 5.10<sup>-5</sup> | 45 to 95 | Membrane of the nuclear transport 2 (NFT2) |
|  | Subtilase | 2.10<sup>-5</sup> to 3.10<sup>-5</sup> | 48 | Hypothetical *C. burnetii* protein with a putative Rif-1 domain |
|  | Rhomboid-1 | 6.10<sup>-4</sup> | 61 | Hypothetical *C. burnetii* protein with a putative OmpA domain |
|  | HtrA | 2e-112 | 100 | *C. burnetii* HtrA protein |

Thus, the only protein that met our screening criteria was the *C. burnetii* (Cb) HtrA-like molecule (CbHtrA). Interestingly, the 4 strains of *C. burnetii* were found to express this hypothetical protein showing 42% identity and 100% coverage with the bacterial HtrA protein of *Salmonella typhimurium* (e value: 2e^−112^) (**Table III)**, representing the best match among the studied sheddases. Among the primary hits, the *in-silico* analysis indicated that the CbHtrA was one of three hits containing a signal peptide compatible with secretion.

### Identification of an HtrA-like sheddase produced by *Coxiella burnetii*

The search for the protein sequence identity of HtrA from *Salmonella typhimurium* in *C. burnetii* revealed that the four strains of *C. burnetii* express a hypothetical protein showing 42% sequence identity and 100% coverage with HtrA (e value: 2e-112).HtrA serine proteases are frequently found in bacteria and have already been described as involved in the catalytic cleavage of E-cad during the human host invasion. These molecules are located in the periplasm and have both protease and chaperon functions. The protease activity is turned on to its active form by heat shock.

The Clustal Omega multiple sequence alignment of 4 strains of *C. burnetiid* – the Cb175 strain, the 3262 strain, the RSA 493 strain and the Z3055 strain – demonstrates that the hypothetical CbHtrA sequence of 451 amino acid residues is 100% identical in the 4 bacterial strains (**Figure 1A**). This hypothetical CbHtrA is characterized by a common organization that includes a signal peptide, a serine protease domain and 2 PDZ (Postsynaptic density protein 95, Drosophila disc large tumor suppressor and Zonula occludens-1 protein domain) domains. CbHtrA exhibits high sequence identity with the HtrA from other bacterial species such as *Tropheryma whipplei*, *Campilobacter jejuni*, *Yersinia pestis*, *Salmonella typhimurium*, Enteropathogenic *Escherichia Coli*, and *Shigella flexneri*, while this sequence is genetically more distant from the HtrA sequences of Mycobacteria and from the HtrA sequences of eukaryotes characterized by a single PDZ domain (**Figure 1B**). Within the serine protease domain, the catalytic triad H, D, S described as essential to function, appears conserved in the hypothetical CbHtrA. Altogether, these results suggest that a secreted form of the CbHtrA protein could be functionally active.

**Figure 1:**
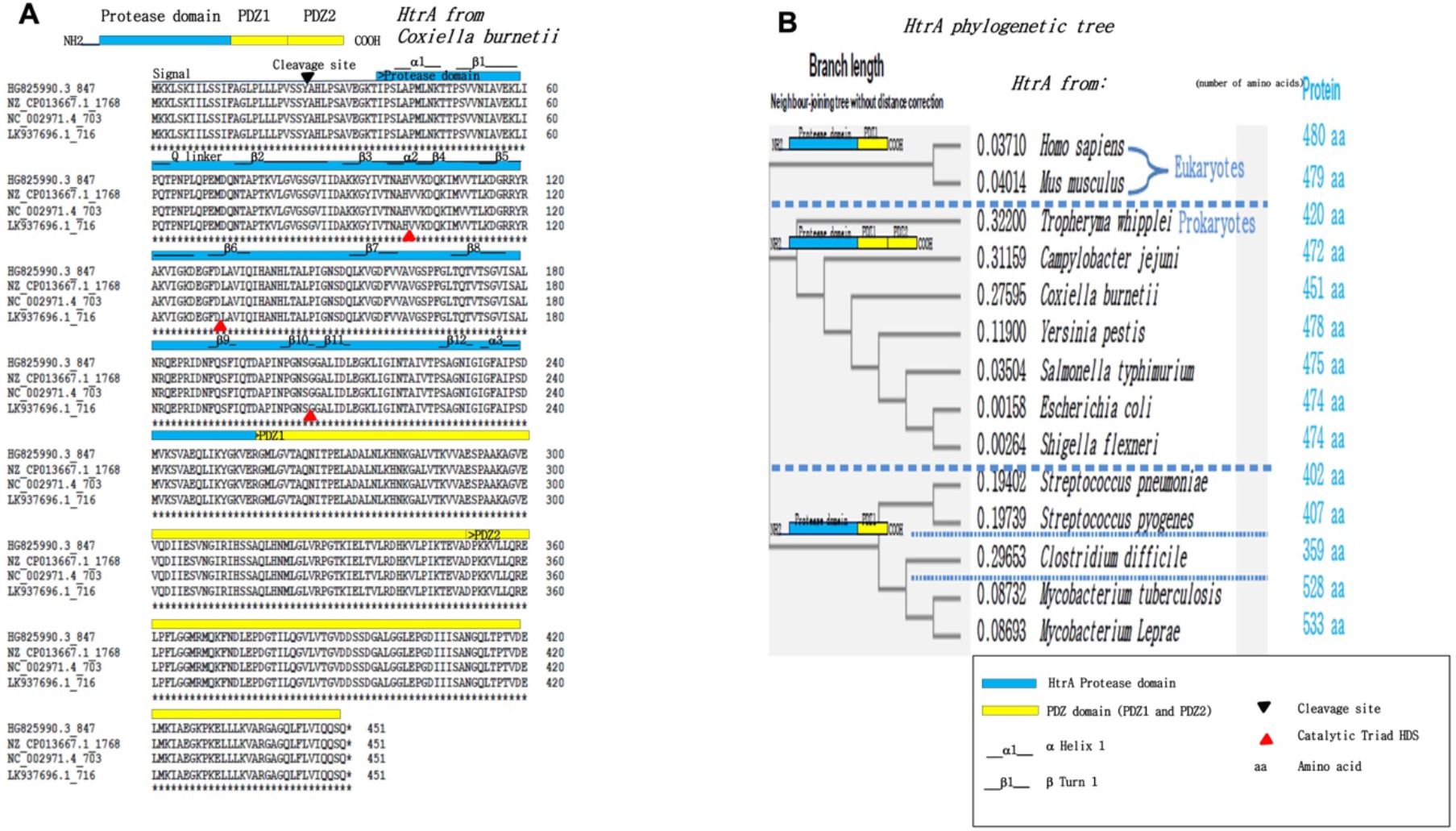
Characterization of heat-shock-induced serine protease HtrA from *Coxiella burnetii*. (A) CLUSTAL sequence alignments of the cbHtrA hypothetical protein from *C. burnetii* Cb175 Guiana strain (Accession nb: HG825990.3), NL3262 Netherland strain (Accession nb: NZ_CP013667.1), RSA 493 Montana USA strain Nine Mile (Accession nb: NC_002971.4), and Z3055 strain (Accession nb: LK937696). The cbHtrA hypothetical protein in the different strains of *C. burnetii* contains 451 amino acid residues and show 100% amino acid identity from one strain to another. The assignment of signal peptide, serine protease domain (that includes the catalytic triad HDS) and PDZ (protein-protein interaction) domains is, according to (81), the catalytic triad H, D, S described as essential to function (33, 51) and appears to be conserved in the hypothetical CbHtrA. (B) The HtrA phylogenetic neighbor-joining tree was built using the clustal Omega program and the proteins’ best match for known HtrA from eukaryotes (human and mouse) and prokaryotes. The length of HtrA proteases varies from one species to another (from 359 amino acids to 533 amino acids). Some prokaryotic HtrA contain two PDZ domains while others only contain one PDZ domain. The proteins’ best match to CbHtrA is with a group of HtrA proteases from prokaryotes including *Campylobacter jejuni*, *Yersinia pestis*, *Salmonella typhimurium* and *Escherichia coli*.

### Transcription of the CbHtrA gene-Like in *C. burnetii* infected cells

The human BeWo cells were previously reported to be susceptible to *C. burnetii* infection and to express high amounts of E-cad at their surface(31),which makes this cell line an interesting and practical cellular model for studying the expression of the CbHtrA gene-Like in these cells once infected with *C. burnetii*. The presence of the CbHtrA gene in the genome of the Nine Mile (NM) and Guiana (Gui) strains of *C. burnetii* was demonstrated by PCR analysis using CbHtrA oligonucleotides (**Figure S1**). A similar PCR amplification on BeWo cells using the same oligonucleotide primers pair demonstrated the lack of amplification of the human HtrA (HuHtrA), confirming the specificity of these primers for the bacterial CbHtrA. Next, the CbHtrA mRNA expression was analyzed by qRT-PCR performed on bacterium-free BeWo cells or BeWo cells infected with *C. burnetii* and evaluated using the primers specific to the bacterial CbHtrA. As shown in **Figure 2A**, a CbHtrA mRNA was expressed in BeWo cells infected either by the live Nine Mile (NM) or Guiana (Gui) strains of *C. burnetii*. It is worth noting that the CbHtrA expression (**Figure 2B**) was significantly higher in the avirulent phase II bacteria than in the virulent phase I of the NM and Gui strains of *C. burnetii*. This result indicates that the CbHtrA mRNA is expressed in *C. burnetii* infected cells and suggests an association between CbHtrA expression and virulence.

**Figure 2.**
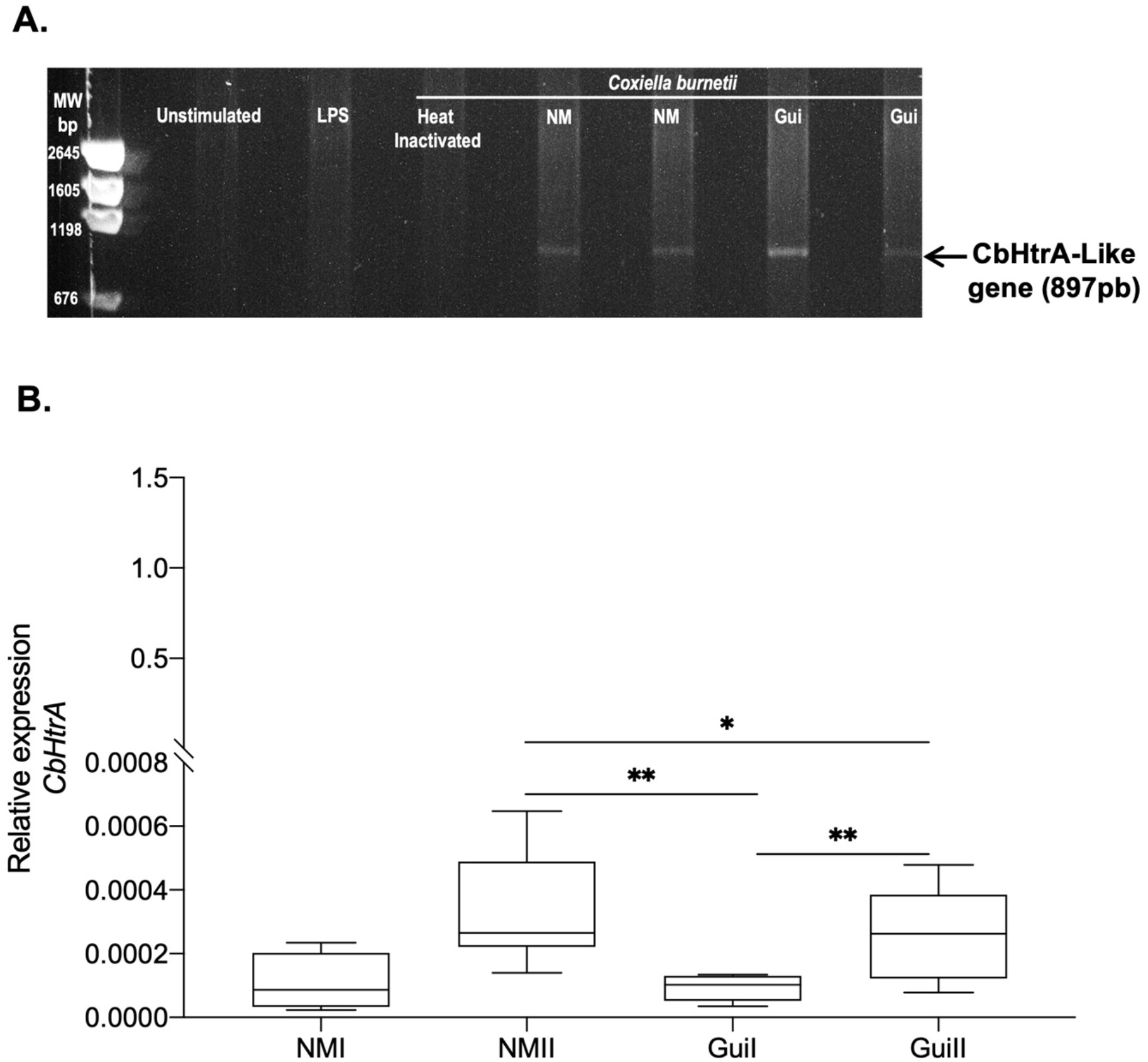
Expression of CbHtrA in BeWo cells infected by *C. burnetii* (n=6). (**A**) RT-PCR amplification on BeWo cells uninfected or infected with the bacterium. Heat inactivated *C. burnetii*, *E. Coli* LPS or unstimulated BeWo cells were used as controls to demonstrate the lack of expression of the *CbHtrA* using the oligonuclotide primers specific to the *CbHtrA*-like gene. (**B**) The qRT-PCR analysis of the *CbHtrA* gene expression in BeWo cells infected by the Nine Mile (NM) and the more aggressive Guiana (Gui) strain of *C. burnetii* using primers specific to the CbHtrA. The expression level of investigated genes was illustrated as ± SEM of the Relative Expression (RE = 2^−ΔCT^, where Δct = [Ct_target_ – Ct_actin_]) and compared using the Mann-Whitney U test. For p value <0.05: symbol *; p value <0.01: symbol **.

### Effects of CbHtrA from *C. burnetii* on E-cadherin expression in BeWo cells

Due to the fact that both CbHtrA and HuHtrA mRNAs can likely be produced in BeWo cells infected by *C. burnetii*, further proof that CbHtrA was functional and competent for cleavage of E-cad was needed. To further explore the functional properties of CbHtrA we designed a synthetic Coxb-HtrA gene mimicking the CbHtrA sequence. The synthetic gene was cloned in the pET-22b (+) bacterial vector optimized for expression in the electrocompetent BL21(DE3) *E. coli* strain. The expressed protein was purified using an ÄKTA avant system equipped with a StrepTrap HP column. Protein expression was assessed by SDS-PAGE that confirmed the presence of a protein migrating at the expected molecular weight (about 50 kDa) and this was confirmed by performing MALDI-TOF MS analysis on previously obtained gel bands (**Figure 3A**). The purified recombinant protein was then tested to evaluate its protease activity. BeWo cells were incubated for 24 hours with recombinant CoxbHtrA protein at different dilutions (1:10; 1:100; 1:1000) from the concentration of 1.6 mg/mL, and the cleavage of E-cad was quantified by SDS-PAGE and immunoblot. In cell lysate from BeWo cells exposed to CoxbHtrA treatment, there was an increase in a 60 kDa proteolytic fragment of E-cad (**Figure 3B**). Unexpectedly, we observed a parallel over-production in the full-length form of E-cad. Also, the same cleavage profile was observed in BeWo cells infected with the bacterium. To further explore these results, the expression level of E-cad protein on the surface of BeWo cells exposed to the CoxbHtrA protein was measured using an antibody targeting the ectodomain portion of E-cad. Thus, the percentage of BeWo cells expressing membrane-bound E-cad was measured and a significant decrease in E-cad expression was found in the CoxbHtrA protein stimulated cells compared to the unstimulated cells used as controls or cells exposed to the elution buffer of the recombinant protein (**Figure 3C**). Furthermore, the culture supernatant of BeWo cells exposed to CoxbHtrA treatment was tested by ELISA for the presence of sE-cad released from cells. Live *C. burnetii* induced a higher level of sE-cad than unstimulated cells, cells stimulated with *E. coli* LPS or cells exposed to heat-inactivated *C. burnetii* (**Figure 3D**). In addition, when cells were exposed to CoxbHtrA, the sE-cad concentration in cell culture supernatants was significantly higher than in all other experimental conditions with a dose-dependent manner, indicating that CoxbHtrA is a functional sheddase.

**Figure 3.**
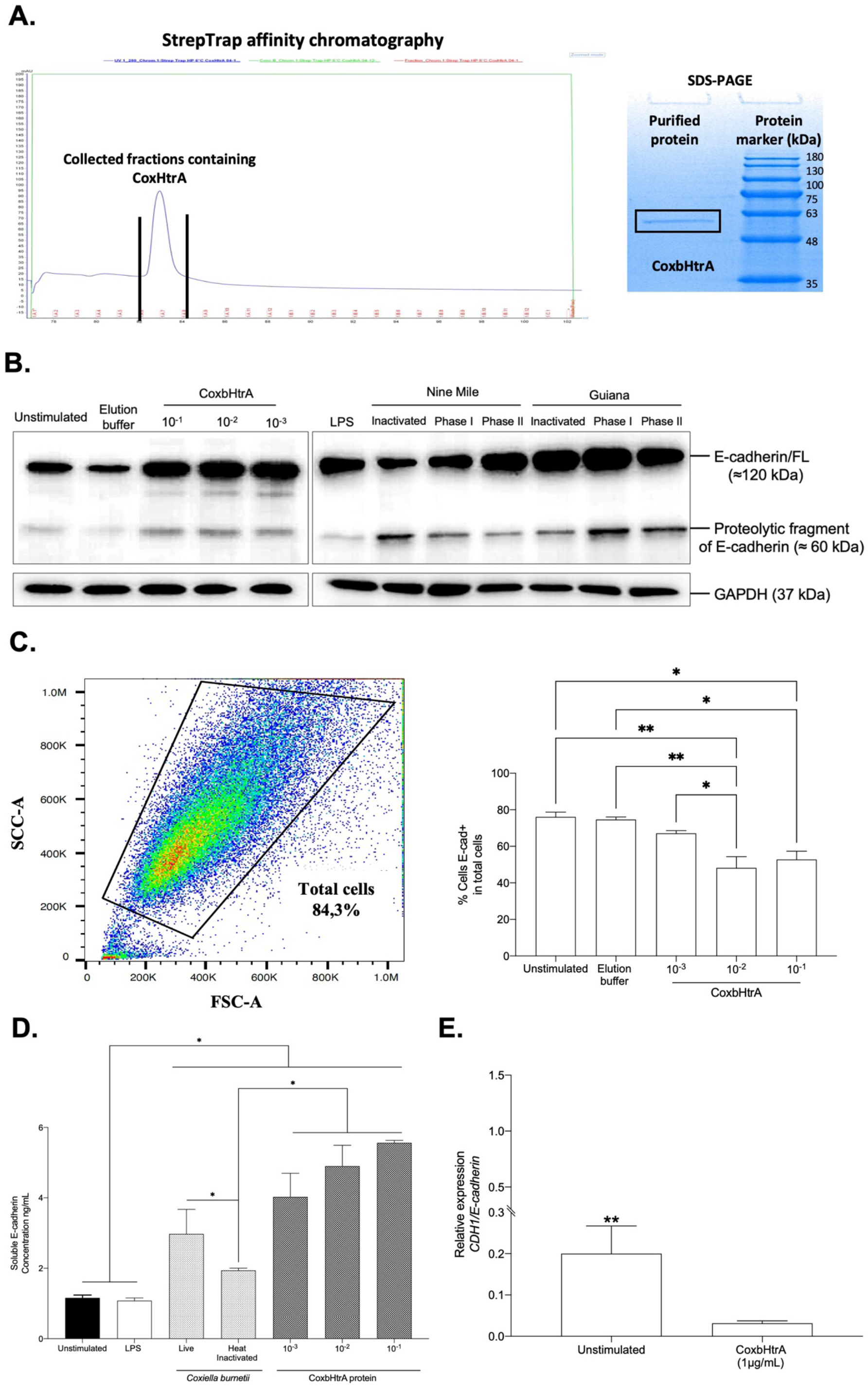
E-cad modulation following a cellular infection by *C. burnetii* or exposure to the recombinant CoxbHtrA protein. (**A**) The tagged recombinant CoxbHtrA protein was purified from transformed *E. coli*. MALDI-TOF (left panel) SDS-PAGE (right panel). (**B**) The quantification by immunoblot of E-cad expression and GAPDH (control) in protein lysates of BeWo cells stimulated with different dilutions of the CoxbHtrA protein and the elution buffer as control. Also, BeWo stimulated with *E. coli* LPS cells or infected with *C. burnetii* NMI laboratory strain and Guiana strain. (**C**) Flow cytometry analysis of E-cad expression at the surface of BeWo cells (n=3). Cells were incubated with recombinant CoxbHtrA (dilution 1:10; 1:100 and 1:1000) or were maintained in a cultured medium without additives (unstimulated or incubated with an elution buffer of the recombinant protein at 1:10 dilution was used as a control). The histogram indicates the percentage of cells expressing E-cad under the different experimental conditions. (**D**) The ELISA quantification of sE-cad released in the supernatant of BeWo cells under different experimental conditions (n=3, unstimulated, *E. coli* LPS-stimulated, inactivated, or live *C. burnetii* Nine Mile strain-infected). (**E**) Relative expression (RE = 2^−ΔCt^) where ΔCt = Ct_Target_ – Ct _Actin_ of CDH1/E-Cad mRNA in BeWo cells untreated or treated with the CoxbHtrA recombinant protein. The non-parametric Mann-Whitney test was used for statistical analysis of all data. For p value < 0.05: symbol *; p value < 0.01: symbol **.

These results indicate that *C. burnetii* secretes its own HtrA and that this sheddase is able to generate a cleavage of the integral E-cad. Moreover, incubation of BeWo cells with CoxbHtrA results in a significant decrease of the CDH1/E-cad gene expression (**Figure 3E**), suggesting the possible activation of a negative feedback loop on the CDH1/E-cad gene expression.

### *C. burnetii* infection of BeWo cells also modulates the expression of human sheddases

Since we had demonstrated the capacity of CoxbHtrA to function as a sheddase with an ability to cleave the integral membrane E-cad, and because several human sheddases (including members of the MMP and ADAM family of sheddases) were previously reported in the literature as being overexpressed in patients with Q fever disease, it was tempting to question whether or not *C. burnetii* could modulate the activity of human sheddases.

First, the HuHtrA mRNA expression was evaluated using another set of primers specific to the Human HtrA. As shown in **Figure 4A**, the expression of HuHtrA mRNA was significantly increased in all infected conditions, compared to unstimulated and LPS stimulated cells, suggesting that the infectious process, besides being associated with the production of CbHtrA, also induces HuHtrA gene expression. It should be emphasized that HuHtrA was also found to be induced when BeWo cells were exposed to heat-inactivated *C. burnetii* and when the cells were incubated with the CoxbHtrA recombinant protein, indicating that infection is not required to activate the expression of HuHtrA but is achieved under bacterial antigen contact or cleavage of E-cad. As shown in **Figure 4B**, when BeWo cells are infected with *C. burnetii*, the immunoblotting detection of HuHtrA molecule provides evidence that a new isoform of HuHtrA called HuHtrA/S is overproduced by the BeWo cells. The expression of human sheddases MMP-3, MMP-9, MMP-12, ADAM-8, ADAM-10 and ADAM-15 in BeWo cells infected by *C. burnetii* was also studied. As shown in **Figure 4C**, we found that the virulent (phase I) Nine Mile strain induces MMP-3, MMP-9 expression and, at a lower level, an expression of ADAM-10 and ADAM-15. This expression seems to be associated with the virulence since infection with the avirulent (phase II) Nine Mile does not induce a similar activation of human sheddase gene expression.

**Figure 4.**
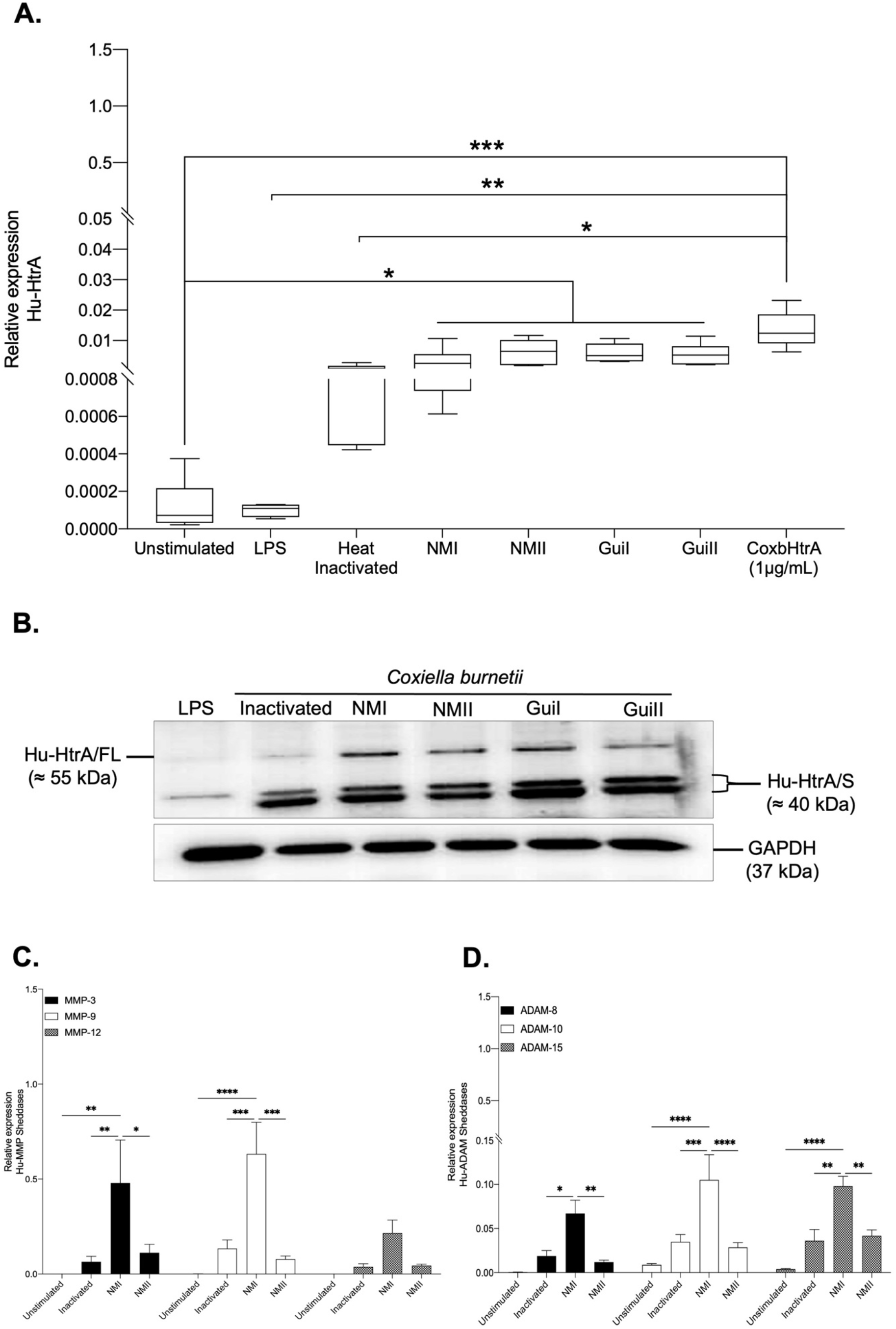
Expression of human sheddases in BeWo cells infected by *C. burnetii*. (**A**) The expression of HuHtrA mRNA in BeWo cells exposed to LPS from *E. coli* and heat-inactivated Guiana *C. burnetii* strain, and BeWo cells infected by either phase I or phase II of Nine Mile or Guiana strain of *C. burnetii* was evaluated by qRT-PCR using primers specific to the HuHtrA. (**B**) Quantification by immunoblot of HuHtrA expression and GAPDH (control) in protein lysates of BeWo cells. The HuHtrA expression was analyzed using an anti-HuHtrA antibody for different experimental conditions in which BeWo cells were incubated with different dilutions of CoxbHtrA protein or the elution buffer as a control (left panel). The HuHtrA was also analyzed for BeWo cells incubated with E. coli LPS or for cells infected with either phase I or phase II of *C. burnetii* NM and Guiana strains (right panel). (**C**) Expression of human sheddases MMP-3, MMP-9, MMP-12, ADAM-8, ADAM-10 and ADAM-15 in BeWo cells infected by *C. burnetii* (n=6). The analysis of human sheddase gene expressions in BeWo cells infected by the Nine Mile strain of *C. burnetii* either in the virulent phase I (NMI) or avirulent phase II (NMII) was performed using qRT-PCR. The control consisted of bacteria-free BeWo cells and BeWo cells exposed to heat-inactivated NM bacteria. The expression level of investigated genes was illustrated as ± SEM of the Relative Expression (RE = 2^−ΔCt^) where ΔCt = Ct_Target_ − Ct _Actin_ and compared using the Mann-Whitney U test. For p value <0.05: symbol *; p value <0.01: symbol **; p value <0.001: symbol ***; p value <0.0001: symbol ****.

These results indicate that during infection of BeWo cells by *C. burnetii*, both prokaryotic (CbHtrA) and eukaryotic (HuHtrA) sheddases are induced with a likely dominance of CbHtrA expression during the avirulent phase II.

### Evaluation of the role of CbHtrA in the physiopathology of *C. burnetii* infection

Although the trophoblastic BeWo cell *in vitro* model was well suited for the functional characterization of CbHtrA and CoxbHtrA, it cannot be considered relevant for investigating the pathophysiology of *C. burnetii* infections. Indeed, monocytes and macrophages are the main targets for *C. burnetii* during infection of humans with the bacteria. More precisely, it was previously reported that monocytes in which *C. burnetii* survives without replication are the classically activated macrophages (CAMs or M1) which exhibit a proinflammatory M1-type response (e.g. TNF, IL-1β, IL-6, IL-12, NO production), whereas macrophages in which *C. burnetii* slowly replicates are polarized towards an anti-inflammatory M2-type response(47) (e.g. TGF-β1, IL-4, IL-6, IL-10, Arginase 1). To study the existence of a possible association between the production of CbHtrA and an effect on the cellular targets of the bacterium, a model of THP-1 cells – differentiated into macrophages which express high levels of E-cad at their surface (**Figure 5A**) and can be induced towards an M1 or M2 polarization – were exposed to the active CoxbHtrA sheddase and analyzed for an M1/M2 mRNA interleukins signature. To further confirm the functional potential of CoxbHtrA, THP-1-macrophages (M0, M1, or M2 types) were analyzed for cell-surface E-cad expression. As shown in **Figure 5B** the population expressing the lowest levels of E-cad were the M2-like type cells, suggesting an association between down regulation E-cad (either at the transcriptional level, protein turn over or shedding) with M2 type differentiation. Next, THP-1 cells polarized into M1 type or M2 type were tested for the expression of several genes either in the presence or absence of active CoxbHtrA sheddase. As shown in **Figures 5C** and **6D**, treatment with CoxbHtrA triggers a significant down-regulation of most genes including IL-6, which is a representative cytokine of the M1 phenotype, and a significant up-regulation of genes such as the IL-10, the expression of which defines the M2 type cells.

**Figure 5.**
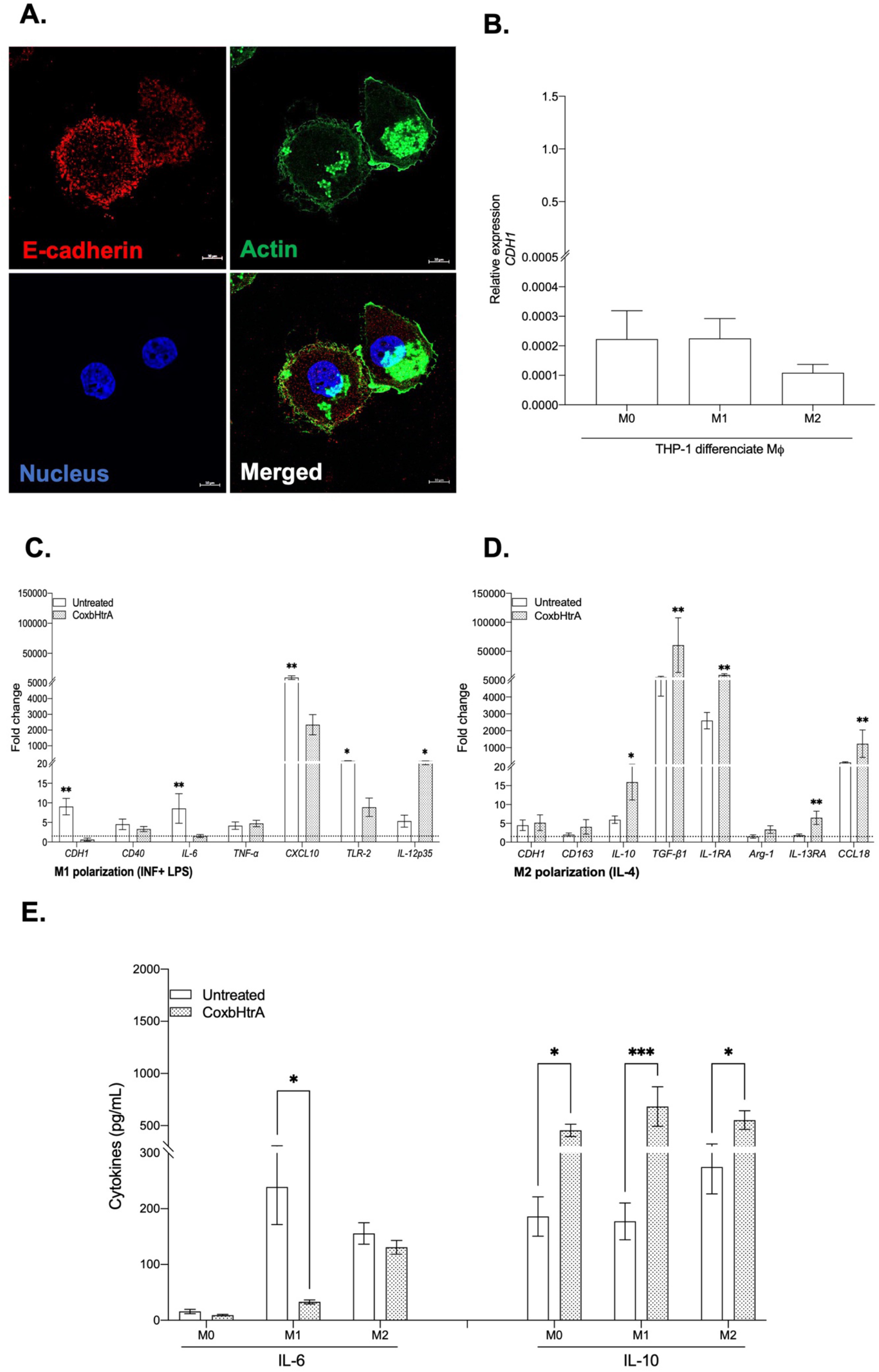
Expression of human E-cad in macrophage-differentiated THP-1. (A) Illustration of single plane confocal microscope analysis of E-cad in THP-1-macrophage. Actin expression was shown as the control as well as the labeling of the nucleus. (B) CDH1/E-cad mRNA expression in THP1 macrophages polarized by treatment with IFN-γ and lipopolysaccharide for the M1 phenotype, IL-4 for the M2 phenotype or without agonist for the M0 phenotype. The results (n=6) are expressed as RE where RE = 2^−ΔCt^. (C) and (D) mRNA expression patterns of different cytokines in THP-1 macrophages (M1 and M2 phenotypes) untreated or treated with CoxbHtrA for 24 hours. The results (n=6) are expressed as Fold Change (FC = 2^(−ΔΔCt)^ where ΔΔCt = (Ct_Target_ − Ct_Actin_)treated − (Ct_Target_ − Ct_Actin_)untreated. Gene expression was considered modulated when the fold change was ≥1.5 (indicated by the dotted line). (E) The release of IL-6 and IL-10 by THP-1 macrophages (n=3) polarized into M1 or M2 phenotypes or unpolarized (M0), and untreated or treated with CoxHtrA, was determined by immunoassay. The data represent a mean ± standard error. Statistical analyses were performed using the Mann-Whitney U test (untreated vs. CoxbHtrA treatment). For p value <0.05: symbol *; p value <0.01: symbol **; p value <0.001: symbol ***.

Since previous publications reported that *C. burnetii* replication occurs in monocytes from patients with Q fever endocarditis who overproduce the IL-10 immunoregulatory cytokine that enables the bacteria to survive in ‘disarmed’ monocytes(48, 49), we aimed to test what the impact of active CoxbHtrA sheddase treatment on the replication of virulent *C. burnetii* could be. Under CoxbHtrA treatment, the bacterial DNA copy number is significantly increased on day 3 and day 6 post-infection compared to untreated cells (**Figure 6A**), suggesting that the CoxbHtrA treatment ‘disarms’ the cells, allowing the bacteria to bypass the latency phase seen in the control experiment. This is consistent with the observation that 24 hours after CoxbHtrA treatment the cells infected with the virulent *C. burnetii* show an increased transcription of the IL-10 mRNA and other mRNA such as TGFβ, indicating a preferential M2 signature (**Figure S2**). This overexpression of IL-10 mRNA is confirmed at the protein level (**Figure 6B**). Moreover, as expected, this IL-10 production is associated with a down-regulation of E-cad expression (**Figure S3**).

**Figure 6.**
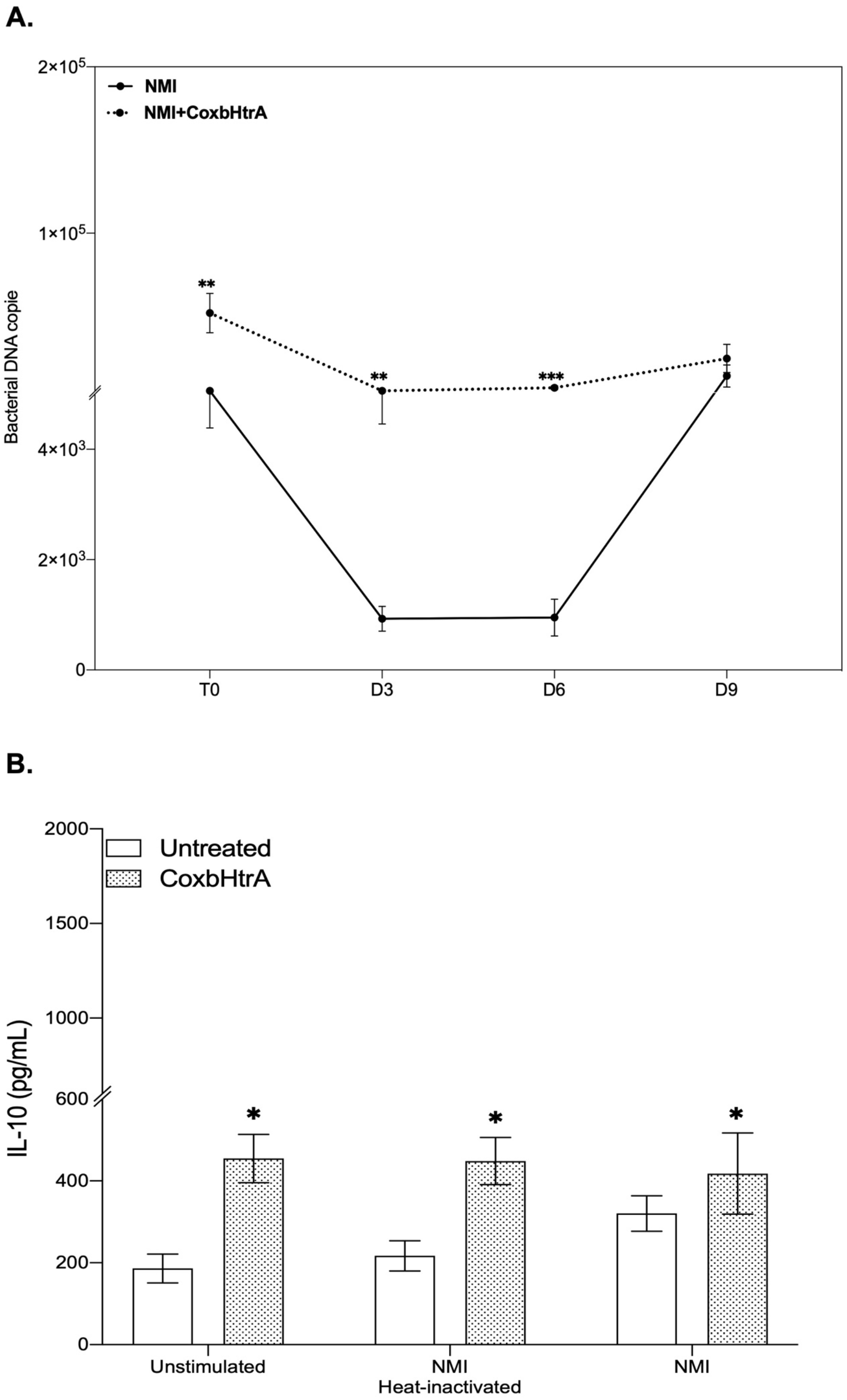
Role of CoxbHtrA in THP-1-macrophages’ response to *C. burnetii*. Macrophages differentiated from THP-1 cells were untreated or treated with CoxbHtrA for 24 hours before infection with C. burnetii (50 bacteria per cell). (A) For *C. burnetii* persistence, THP-1-macrophage cells (n=8) were infected for 4 hours – a period designed as the uptake phase (or T0). After the bacterial uptake phase, the cells were cultured for 9 days, and the presence of bacterial DNA copies was assessed every 3 days by qPCR (in duplicate). (B) The release of IL-10 by THP-1 macrophage (n=3) with or without pretreatment and infected by live *C. burnetii* or exposed to heat-inactivated bacteria for 24 hours was determined by immunoassays. Data represent mean ± standard error. Statistical analyses were performed using the Mann-Whitney U test (untreated vs. CoxbHtrA treatment). For p value <0.05: symbol *; p value <0.01: symbol **; p value <0.001: symbol ***.

## Discussion

So far, many results previously published in the literature argued in favor of an activation of MMP metalloproteinase and ADAM family of human sheddases during *C. burnetii* infections. The activation of such sheddases in human cells could provide a rational hypothesis to explain why we found high concentrations of sE-cad in sera of Q fever patients, and why infection of BeWo cells by *C. burnetii* leads to the modulation of the E-cad/β-cat pathway(27, 31). However, apart from the human sheddases that were likely to contribute to E-cad proteolysis and sE-cad release in Q fever patients, we also wanted to explore the possibility of a direct mechanism of cleavage initiated by the bacterium itself, assuming the possible synthesis of a sheddase encoded in the genome of *C. burnetii*.

Several bacteria have previously been found to be able to encode sheddases that cleave E-cad(28). In light of this observation, it was possible to postulate that *C. burnetii*, like many other bacteria species, encodes a sheddase or sheddases in its genome. To improve our knowledge on putative sheddases produced by *C. burnetii* itself, we used *in silico* analysis for an initial screening. Among the hits we found hypothetical proteins named CbMMP-3-like and CbMMP-9-like presenting short sequence identity with HuMMP3 and HuMMP9. The blasting search in the NCBI database revealed that the CbMMP-3-like sequence did indeed correspond to the nuclear transport factor 2 protein and that the CbMMP-9-like sequence corresponded to the L28 protein of the large 50S ribosomal subunit of *C. burnetii*. In bacteria, the L28 protein has been shown to be encoded by the rpmB operon and required for ribosome assembly in association with the L33 protein(50). Because the nuclear transport factor 2 protein showed unexpected similarities with HuMMP-3 in the region that correspond to the catalytic domain of HuMMP-3 and shared some similarities with the BFT toxin bacterial sheddase, we performed a reverse BLAST that led us to the conclusion that it is very unlikely that such a molecule might exhibit sheddase activity. Using the same approach, we also eliminated the L28 protein and the CbADAM-15 like as possible candidate for sheddase activity. Two additional hypothetical *C. burnetii* proteins with either Rif1 or OmpA domain emerged from the *in silico* screening. However, none of those molecules met the criteria for a possible sheddase. Among the bacterial sheddases used in this study to search hits in *C. burnetii* only one, HtrA(32), turned out to be of high interest because the CbHtrA-like sequence of RSA 493 Nine Mile, NL3262 Netherland, and Z3055 strains were identical over the 451 amino acid residues and because of the probable capacity for secretion. The hypothetical CbHtrA exhibited high sequence identity with the HtrA from bacteria such as *S. typhimurium*, *Enteropathogenic E. Coli*, and *S. flexneri* with a conserved secondary structure organization that included a signal peptide, a serine protease domain and two PDZ domains. Within the serine protease domain, the catalytic triad H, D, S(33, 51) was conserved. To determine if this protein could be responsible for the cleavage of E-cad during *C. burnetii* infection, cells were infected *in vitro* with *C. burnetii* and the expression of the CbHtrA gene was evaluated, which turned out to be expressed. Furthermore, a recombinant CoxbHtrA protein was found to act as a sheddase as it was able to trigger sE-Cad shedding when incubated with BeWo cells that express E-Cad at their surface.

Yet many questions remain to be addressed. The purified CoxbHtrA protein used in our study was obtained after lysis of competent BL21(DE3) expressing the recombinant protein. Therefore, the question must be asked whether this is physiological and whether the CbHtrA sheddase produced by the *C. burnetii* can actually be produced in the extracellular environment to cleave E-cad on the cell surface. *C. burnetii* is a gram negative strict intracellular bacterium, therefore the mechanism by which this protease cleaves the E-cad expressed at the cell membrane remains obscure. Protein trafficking across the bacterial envelope is a complex process that can possibly influence the signaling and integrity of the host cell. The bacterium uses a number of nanomachines (protein secretion system of various types) to allow protein trafficking and export across membranes(52). Pathogenic gram-negative bacteria secrete effector proteins into the periplasm, outer membrane or external milieu by different secretion pathways(53, 54). In light of the other bacterial models in which the bacterium produces an HtrA, the hypothesis of secretion by transport vesicles could be put forward. Similarly, it is possible to imagine that in the bacterial population, certain bacteria could lyse and thus directly release the soluble HtrA into the extracellular environment. In gram-negative bacteria (including *Shigella* and pathogenic *E. Coli*), there is a growing family of serine proteases secreted to the external milieu by a secretion mechanism called an autotransporter pathway(55, 56). In the model of *C. jejuni* it was demonstrated that HtrA is expressed in the periplasmic space, but it can also be secreted into the extracellular environment and cleave the E-cad adherens junctions(57–59). HtrAs form oligomers and these proteases are present in two functional states: a resting state (inactive) and an active state whose tertiary and quaternary structures differ(60). Assembly into oligomers is transient and the proteins return to the resting state as soon as the substrate is depleted(61, 62). Interestingly, *Chlamydia trachomatis*, an obligate intracellular human pathogen as *C. burnetii*, produces HtrA in the periplasm through a sec-dependent pathway. HtrA is also exported outside via an outer membrane vesicle mechanism and is found in the host cell cytosol(63–65). In addition, it was reported that *Glaesserella* (*Haemophilus*) *parasuis* infection activates the canonical Wnt/β-catenin signaling pathway leading to the disruption of the epithelial barrier, with a sharp degradation of E-cadherin and an increase of the epithelial cell monolayer permeability(66), which is an observation very similar to what we recently reported with *C. burnetii*(31).

We have also evidenced that *C. burnetii* infection induces HuHtrA and that the level of induction differs with respect to the virulent phase I or avirulent phase II. So far, it is not clear why such variations are observed or what the respective influence of CbHtrA and HuHtrA is on the cleavage of E-cad during *C. burnetii* infection. In this paper, we have also studied the activation of MMP (MMP-3, MMP-9, MMP-12) and ADAM (ADAM-8, ADAM-10 and ADAM-15) gene expression in BeWo cells infected by *C. burnetii*. We found that the virulent (phase I) Nine Mile strain induces MMP-3, MMP-9 expression and, at a lower level, an expression of ADAM-10 and ADAM-15. Infection with the avirulent (phase II) Nine Mile do not show such drastic induction of human sheddases. The over-expression of MMP molecules was previously reported in Q fever patients(29) or after cells had been exposed *in vitro* to *C. burnetii*(30). MMP are zinc-dependent proteases(67, 68). The zymogen of MMP-3 (proMMP-3), a 54kDa protein, is activated in the 45kDa and 23kDa active forms by limited proteolysis catalyzed by elastase and cathepsin G(69). The active form of MMP-3 was reported to be over-expressed in breast cancer and found to catalyze E-cad proteolysis in that cancer(70, 71). MMP-3 is an activator of the proMMP-9(72). The proMMP-9 is secreted as a 92kDa. The active MMP-9 enzyme is an 82kDa gelatinase that readily digests gelatin(73). Zona occludens 1, α1-Antiproteinase, latent TGF-β1, latent VEGF, fibrin, and NG2 proteoglycan are also substrates of MMP-9(74). Interestingly, MMP-9 was also identified as a key gene in mantle cell lymphoma(75), a non-Hodgkin lymphoma. An over-expression of human MMP-3 in dendritic cells (DC) infected *in vitro* by *C. burnetii* and variations in the HuMMP-9 expression in human cells exposed *in vitro* to heat-inactivated *C. burnetii* was also observed using a screening of differential expressions of 45,000 genes by microarray analysis (data not shown). It is possible that the over-expression of MMP-3 modulates MMP-9 expression in cells infected by *C. burnetii*(72), thereby contributing to the physiopathology of persistent Q fever. MMP-9 over-expression was previously reported following bacterial lipopolysaccharide (LPS) stimulation of lung alveolar macrophages(76). In addition, an over-expression of human HuADAM-15 in persistent Q fever patients’ blood cells and variations in the HuADAM-10 expression in human cells exposed *in vitro* to heat-inactivated *C. burnetii* were also observed using our microarray approach. The HuADAM-15 contains a signal peptide, a metalloproteinase domain, followed by a disintegrin-like domain, cysteine-rich domain, epidermal growth factor domain, short connecting linker, hydrophobic transmembrane segment and cytoplasmic tail. The protein contains a functional catalytic consensus sequence (HEXGEHXXGXXH). ADAM-15 has been linked to a number of different cancerous diseases(77) as well as to the modulation of epithelial cell-tumor cell interactions(78). It was reported in the literature that the ectodomain shedding of E-cad by ADAM-15 supports the ErbB receptor activation associated with the progression of prostate and breast cancer(79). Altogether, these data indicate that *C. burnetii* infection is associated with the modulation of several sheddases belonging to the MMPs and/or ADAM families and that qualitative and/or quantitative individual variations can be evidenced from patient to patient during *C. burnetii* infections. It is also worth noting that when human sheddases are highly expressed in association with *C. burnetii* infection (e.g. Nine Mile phase I), the CbHtrA is not highly expressed, while in Nine Mile phase II infection the CbHtrA is highly expressed and the HuMMP and HuADAM are less expressed, suggesting the existence of a molecular mechanism involved in regulating the balance between the prokaryotic and eukaryotic sheddase gene expression.

Intracellular bacteria replication is likely controlled by the levels of IL-10, an immunoregulatory cytokine that is known to inhibit the production of reactive oxygen and reactive nitrogen intermediates, thereby preventing the generation of toxic compounds by these intracellular bacteria(80). It was previously reported that *C. burnetii* replication occurs in monocytes from patients with Q fever endocarditis who overproduce IL-10, while the bacteria are killed in monocytes from patients with acute Q fever, suggesting that IL-10 enables monocytes to support *C. burnetii* replication and favors the development of chronic Q fever(49). Here, we demonstrated that the recombinant active CoxbHtrA sheddase triggers a massive induction of IL-10 in THP-1 cells characteristic of an M2-type response and we found that the lowest levels of E-cad expression were in the M2-like type cells, suggesting an association between down regulation E-cad with M2 type differentiation and the induction of IL-10 production. Under CoxbHtrA treatment, the NMI virulent bacterial DNA copy number is significantly increased on day 3 and day 6 post-infection compared to untreated cells, suggesting that bacterial replication is boosted through the action of the CoxbHtrA sheddase. For the first time, these results partially lift the veil on the mechanism that enables the bacteria to replicate intracellularly.

Although the complete pattern of molecular crosstalk between *C. burnetii* and its target cell leading to the shedding of soluble E-cad could be more complex (with concurrent expression of HuHtrA and likely other human sheddases) than simply a direct action of CbHtrA, our data provide the first direct evidence that *C. burnetii* encodes its own CbHtrA sheddase that are able to cleave E-Cad. This is also the first description of a functional HtrA sheddase encoded in the *C. burnetii* genome of 4 different strains of this bacteria. Our data indicate that *C. burnetii* possesses a proteolysis strategy to manipulate host cell signaling pathways via the secretion of CbHtrA, which may play an important role in interactions with host cells and bacterial virulence. Moreover, this work clearly establishes a link between the presence of gene coding for a functional HtrA sheddase (CbHtrA) in the genome of *C*. *burnetii* with the ability of this gene to encode a functional sheddase (CoxbHtrA) that cleaves the E-cad, leading to an M2 polarization of the target cells and inducing their secretion of IL-10.

## Acknowledgments

We thank Dr. Pierre Pontarotti, Benoit Desnues, Matteo Bonazzi and Muriel Masi for their helpful discussions. We also thank Philippe Decloquement for his expert technical assistance with Maldi-Tof. This work was supported by the French Government under the “Investissements d’avenir” (Investments for the Future) program managed by the “Agence Nationale de la Recherche” (National Research Agency) (reference: Méditerranée Infection 10-IAHU-03 grant to Didier Raoult).

## Author contributions

I.O.O. performed the biological experiments. A.C., C.D. and A.L. performed the bioinformatics analysis. L.P. produced the recombinant CoxbHtrA protein. C.D. and J.L.M supervised the experimental work. I.O.O and C.D. conceived and wrote the first draft of the manuscript. All authors participated in the writing and corrections of the manuscript. All authors read the manuscript and agreed with its content.

## Disclosure and conflict of interest

The authors declare that the research was conducted in the absence of any commercial or financial relationships that could be construed as a potential conflict of interest.

---

**Figure S1.**
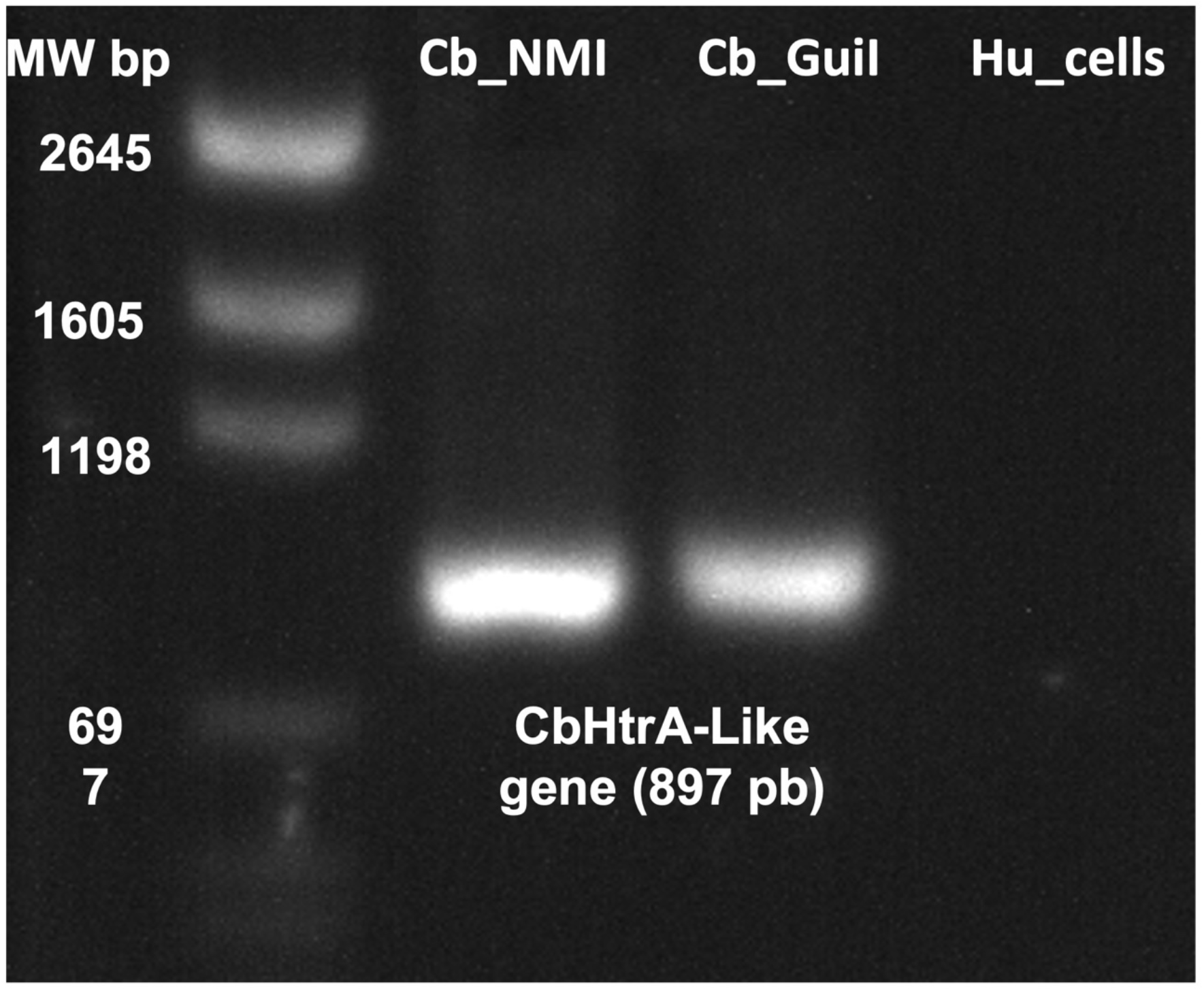
The presence of the CbHtrA gene in the Nine Mile (NM) and Guiana (Gui) strains of *C. burnetii*. Specific amplification by PCR of a fragment of 640 bp corresponding to the size expected using the oligonucleotide primers chosen. This oligonucleotides pair does not amplify the HuHtrA.

**Figure S2.**
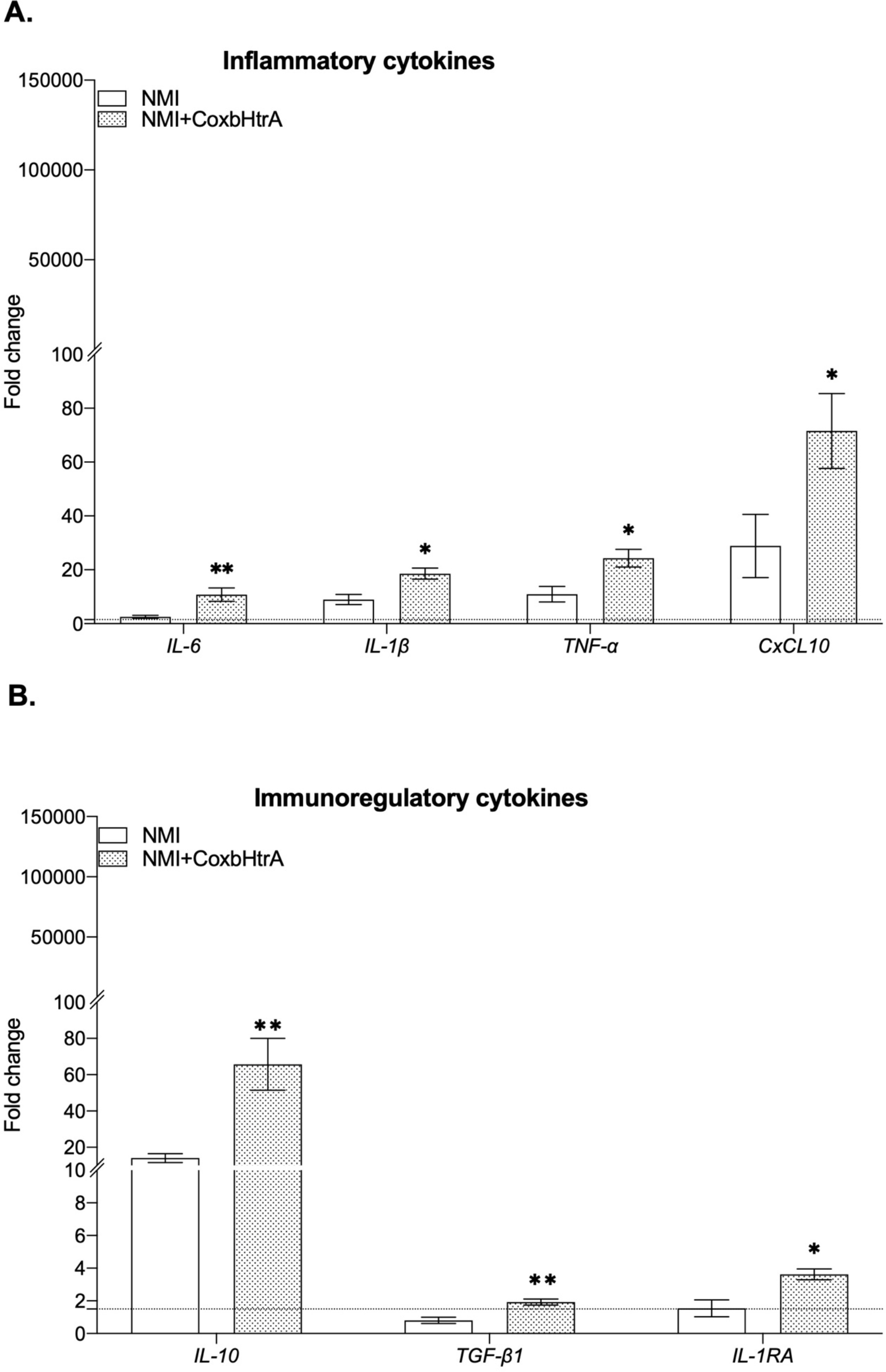
The role of CoxbHtrA in the THP-1-macrophages’ response to C. burnetii. Macrophages differentiated from THP-1 cells were treated with CoxbHtrA for 24 hours before infection with C. burnetii (50 bacteria per cell). The expression of THP-1 macrophages (A) M1 and (B) M2 polarization genes was investigated by qRT-PCR. The results (n=6) are expressed as Fold Change (FC = 2^(−ΔΔCt)^ where ΔΔCt = (Ct_Target_ − Ct_Actin_)treated/infected − (Ct_Target_ − Ct_Actin_)untreated/uninfected. Gene expression was considered modulated when the fold change was ≥ 1.5 (indicated by the dotted line). The data represent mean ± standard error. Statistical analyses were performed using the Mann-Whitney U test (untreated vs. CoxbHtrA treatment). For p value <0.05: symbol *; p value <0.01: symbol **

**Figure S3.**
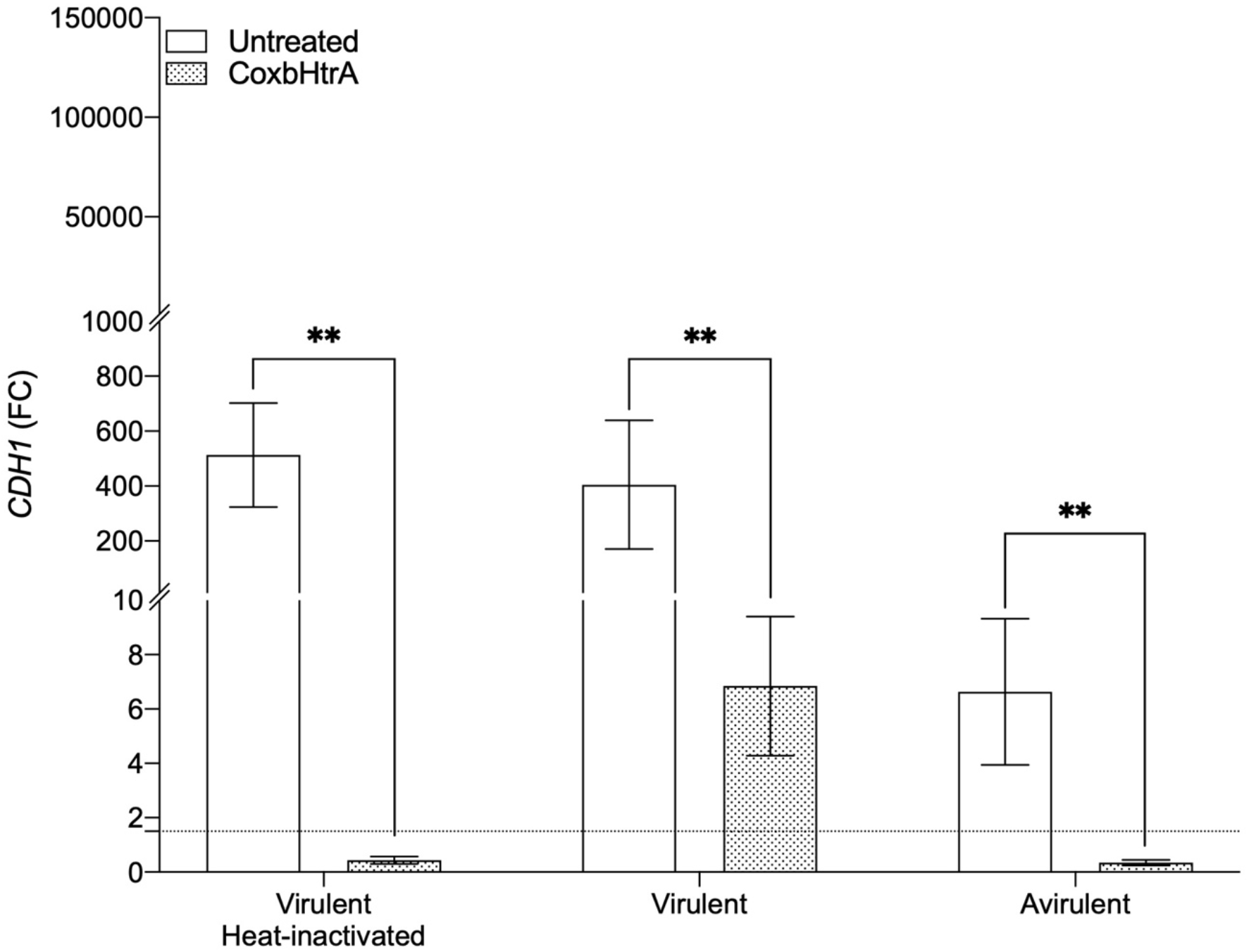
Expression of human CDH1/E-cad mRNA in macrophages differentiated from THP-1 (n=3) with or without pre-treatment, infected by live or heat-inactivated virulent or avirulent *C. burnetii* strains. The results (n=6) are expressed as Fold Change (FC = 2^(−ΔΔCt)^ where ΔΔCt = (Ct_Target_ − Ct_Actin_)treated/infected − (Ct_Target_ − Ct_Actin_)untreated/uninfected. Gene expression was considered modulated when the fold change was ≥ 1.5 (indicated by the dotted line). The data represent mean ± standard error. Statistical analyses were performed using the Mann-Whitney U test (untreated vs. CoxbHtrA treatment). For p value <0.01: symbol **.

## Notes

### Competing Interest Statement

The authors have declared no competing interest.

## References

1. M. Maurin, D. Raoult, Q fever. Clin Microbiol Rev 12, 518–553 (1999).

2. A. O. Barry, J.-L. Mege, E. Ghigo, Hijacked phagosomes and leukocyte activation: an intimate relationship. Journal of Leukocyte Biology 89, 373–382 (2011).

3. J. Pechstein, et al., The Coxiella burnetii T4SS Effector AnkF Is Important for Intracellular Replication. Front Cell Infect Microbiol 10, 559915 (2020).

4. E. E. McClure, et al., Engineering of obligate intracellular bacteria: Progress, challenges and paradigms. Nature Reviews Microbiology 15, 544–558 (2017).

5. E. J. van Schaik, C. Chen, K. Mertens, M. M. Weber, J. E. Samuel, Molecular pathogenesis of the obligate intracellular bacterium Coxiella burnetii. Nat Rev Microbiol 11, 561–573 (2013).

6. C. Melenotte, et al., Clinical Features and Complications of Coxiella burnetii Infections From the French National Reference Center for Q Fever. JAMA Network Open 1, e181580 (2018).

7. C. Melenotte, et al., B-cell non-Hodgkin lymphoma linked to Coxiella burnetii. Blood 127, 113–121 (2016).

8. C. Melenotte, et al., Coxiella burnetii: A Hidden Pathogen in Interstitial Lung Disease. Clin Infect Dis 67, 1120–1124 (2018).

9. C. Melenotte, et al., The hypervirulent Coxiella burnetii Guiana strain compared in silico, in vitro and in vivo to the Nine Mile and the German strain. Clinical Microbiology and Infection 25, 1155.e1–1155.e8 (2019).

10. S. E. van Roeden, P. C. Wever, J. J. Oosterheert, “Chronic Q fever-related complications and mortality: data from a nationwide cohort” - Author’s reply. Clin Microbiol Infect 25, 1436–1437 (2019).

11. J. M. Weehuizen, et al., No increased risk of mature B-cell non-Hodgkin lymphoma after Q fever detected: results from a 16-year ecological analysis of the Dutch population incorporating the 2007-2010 Q fever outbreak. Int J Epidemiol, dyac053 (2022).

12. P. T. G. Elkington, C. M. O’Kane, J. S. Friedland, The paradox of matrix metalloproteinases in infectious disease. Clin Exp Immunol 142, 12–20 (2005).

13. I. Vanlaere, C. Libert, Matrix metalloproteinases as drug targets in infections caused by gram-negative bacteria and in septic shock. Clin Microbiol Rev 22, 224–239, Table of Contents (2009).

14. F. E. Ortega, et al., Adhesion to the host cell surface is sufficient to mediate Listeria monocytogenes entry into epithelial cells. Mol. Biol. Cell 28, 2945–2957 (2017).

15. E. Kague, et al., Methylation status of CDH1 gene in samples of gastric mucous from Brazilian patients with chronic gastritis infected by Helicobacter pylori. Arq Gastroenterol 47, 7–12 (2010).

16. T. Barisani-Asenbauer, et al., Chlamydia trachomatis infection induces epithelial-mesenchymal transition in conjunctival epithelial cells. Investigative Ophthalmology & Visual Science 58, 5778 (2017).

17. G. Jacobs, et al., Polymorphisms in the 3′-untranslated region of the CDH1 gene are a risk factor for primary gastric diffuse large B-cell lymphoma. Haematologica 96, 987–995 (2011).

18. S. Wu, K.-C. Lim, J. Huang, R. F. Saidi, C. L. Sears, Bacteroides fragilis enterotoxin cleaves the zonula adherens protein, E-cadherin. Proc Natl Acad Sci U S A 95, 14979–14984 (1998).

19. L. Chung, et al., Bacteroides fragilis Toxin Coordinates a Pro-carcinogenic Inflammatory Cascade via Targeting of Colonic Epithelial Cells. Cell Host Microbe 23, 203–214.e5 (2018).

20. R. Kumar, et al., Streptococcus gallolyticus subsp. gallolyticus promotes colorectal tumor development. PLOS Pathogens 13, e1006440 (2017).

21. C. M. Niessen, D. Leckband, A. S. Yap, Tissue organization by cadherin adhesion molecules: dynamic molecular and cellular mechanisms of morphogenetic regulation. Physiol Rev 91, 691–731 (2011).

22. F. Hyafil, C. Babinet, F. Jacob, Cell-cell interactions in early embryogenesis: a molecular approach to the role of calcium. Cell 26, 447–454 (1981).

23. B. D. Shields, et al., Loss of E-Cadherin Inhibits CD103 Antitumor Activity and Reduces Checkpoint Blockade Responsiveness in Melanoma. Cancer Res 79, 1113–1123 (2019).

24. P. D. McCrea, M. T. Maher, C. J. Gottardi, Nuclear Signaling from Cadherin Adhesion Complexes. Curr Top Dev Biol 112, 129–196 (2015).

25. M. M. Grabowska, M. L. Day, Soluble E-cadherin: More Than a Symptom of Disease. Front Biosci (Landmark Ed) 17, 1948–1964 (2012).

26. K. N. Syrigos, et al., Circulating soluble E-cadherin levels are of prognostic significance in patients with multiple myeloma. Anticancer Res 24, 2027–2031 (2004).

27. S. Mezouar, et al., High Concentrations of Serum Soluble E-Cadherin in Patients With Q Fever. Front. Cell. Infect. Microbiol. 9, 219 (2019).

28. C. A. Devaux, S. Mezouar, J.-L. Mege, The E-Cadherin Cleavage Associated to Pathogenic Bacteria Infections Can Favor Bacterial Invasion and Transmigration, Dysregulation of the Immune Response and Cancer Induction in Humans. Frontiers in Microbiology 10(2019).

29. L. C. Krajinović, et al., Serum levels of metalloproteinases and their inhibitors during infection with pathogens having integrin receptor-mediated cellular entry. Scand J Infect Dis 44, 663–669 (2012).

30. A. F. M. Jansen, et al., Involvement of matrix metalloproteinases in chronic Q fever. Clin Microbiol Infect 23, 487.e7–487.e13 (2017).

31. I. O. Osman, S. Mezouar, D. Belhaouari-Brahim, J.-L. Mege, C. A. Devaux, Modulation of the E-cadherin/β-catenin signaling pathway in human cells infected in vitro with Coxiella burnetii. 2022.12.08.519566 (2022).

32. C. M. Abfalter, et al., HtrA-mediated E-cadherin cleavage is limited to DegP and DegQ homologs expressed by gram-negative pathogens. Cell Communication and Signaling 14, 30 (2016).

33. M. Israeli, et al., Distinct Contribution of the HtrA Protease and PDZ Domains to Its Function in Stress Resilience and Virulence of Bacillus anthracis. Frontiers in Microbiology 10(2019).

34. R. Toman, L. Skultéty, Structural study on a lipopolysaccharide from Coxiella burnetii strain Nine Mile in avirulent phase II. Carbohydr Res 283, 175–185 (1996).

35. T. A. Hoover, D. W. Culp, M. H. Vodkin, J. C. Williams, H. A. Thompson, Chromosomal DNA deletions explain phenotypic characteristics of two antigenic variants, phase II and RSA 514 (crazy), of the Coxiella burnetii nine mile strain. Infect Immun 70, 6726–6733 (2002).

36. C. Capo, et al., Subversion of monocyte functions by coxiella burnetii: impairment of the cross-talk between alphavbeta3 integrin and CR3. J Immunol 163, 6078–6085 (1999).

37. R. Seshadri, et al., Complete genome sequence of the Q-fever pathogen Coxiella burnetii. Proc Natl Acad Sci U S A 100, 5455–5460 (2003).

38. R. Kuley, et al., First Complete Genome Sequence of the Dutch Veterinary Coxiella burnetii Strain NL3262, Originating from the Largest Global Q Fever Outbreak, and Draft Genome Sequence of Its Epidemiologically Linked Chronic Human Isolate NLhu3345937. Genome Announc 4, e00245–16 (2016).

39. F. D’Amato, et al., The genome of Coxiella burnetii Z3055, a clone linked to the Netherlands Q fever outbreaks, provides evidence for the role of drift in the emergence of epidemic clones. Comp Immunol Microbiol Infect Dis 37, 281–288 (2014).

40. D. Hyatt, et al., Prodigal: prokaryotic gene recognition and translation initiation site identification. BMC Bioinformatics 11, 119 (2010).

41. T. N. Petersen, S. Brunak, G. von Heijne, H. Nielsen, SignalP 4.0: discriminating signal peptides from transmembrane regions. Nat Methods 8, 785–786 (2011).

42. H. Schwende, E. Fitzke, P. Ambs, P. Dieter, Differences in the state of differentiation of THP-1 cells induced by phorbol ester and 1,25-dihydroxyvitamin D _3_. J Leukoc Biol 59, 555–561 (1996).

43. M. Daigneault, J. A. Preston, H. M. Marriott, M. K. B. Whyte, D. H. Dockrell, The Identification of Markers of Macrophage Differentiation in PMA-Stimulated THP-1 Cells and Monocyte-Derived Macrophages. PLoS ONE 5, e8668 (2010).

44. W. Chanput, J. J. Mes, H. F. J. Savelkoul, H. J. Wichers, Characterization of polarized THP-1 macrophages and polarizing ability of LPS and food compounds. Food Funct 4, 266–276 (2013).

45. J. Winer, C. K. Jung, I. Shackel, P. M. Williams, Development and validation of real-time quantitative reverse transcriptase-polymerase chain reaction for monitoring gene expression in cardiac myocytes in vitro. Anal Biochem 270, 41–49 (1999).

46. S. Mezouar, et al., Full-Term Human Placental Macrophages Eliminate Coxiella burnetii Through an IFN-γ Autocrine Loop. Front. Microbiol. 10, 2434 (2019).

47. A. B. Amara, Y. Bechah, J.-L. Mege, “Immune Response and Coxiella burnetii Invasion” in Coxiella Burnetii: Recent Advances and New Perspectives in Research of the Q Fever Bacterium, Advances in Experimental Medicine and Biology., R. Toman, R. A. Heinzen, J. E. Samuel, J.-L. Mege, Eds. (Springer Netherlands, 2012), pp. 287–298.

48. C. Capo, et al., Production of interleukin-10 and transforming growth factor beta by peripheral blood mononuclear cells in Q fever endocarditis. Infect Immun 64, 4143–4147 (1996).

49. E. Ghigo, C. Capo, D. Raoult, J. L. Mege, Interleukin-10 stimulates Coxiella burnetii replication in human monocytes through tumor necrosis factor down-modulation: role in microbicidal defect of Q fever. Infect Immun 69, 2345–2352 (2001).

50. B. A. Maguire, D. G. Wild, The roles of proteins L28 and L33 in the assembly and function of Escherichia coli ribosomes in vivo. Mol Microbiol 23, 237–245 (1997).

51. A. Gieldon, et al., Distinct 3D Architecture and Dynamics of the Human HtrA2(Omi) Protease and Its Mutated Variants. PLoS One 11, e0161526 (2016).

52. A. 2022 Filloux, Bacterial protein secretion systems: Game of types. Microbiology 168, 001193.

53. M. Desvaux, M. Hébraud, R. Talon, I. R. Henderson, Secretion and subcellular localizations of bacterial proteins: a semantic awareness issue. Trends Microbiol 17, 139–145 (2009).

54. J. M. Crane, L. L. Randall, The Sec System: Protein Export in Escherichia coli. EcoSal Plus 7(2017).

55. F. Ruiz-Perez, J. P. Nataro, Bacterial serine proteases secreted by the autotransporter pathway: classification, specificity, and role in virulence. Cell Mol Life Sci 71, 745–770 (2014).

56. K. R. Clarke, et al., Phylogenetic Classification and Functional Review of Autotransporters. Front Immunol 13, 921272 (2022).

57. B. Hoy, et al., Distinct roles of secreted HtrA proteases from gram-negative pathogens in cleaving the junctional protein and tumor suppressor E-cadherin. J Biol Chem 287, 10115–10120 (2012).

58. M. Boehm, et al., Extracellular secretion of protease HtrA from Campylobacter jejuni is highly efficient and independent of its protease activity and flagellum. Eur J Microbiol Immunol (Bp) 3, 163–173 (2013).

59. M. Neddermann, S. Backert, Quantification of serine protease HtrA molecules secreted by the foodborne pathogen Campylobacter jejuni. Gut Pathogens 11, 14 (2019).

60. Z. Chang, The function of the DegP (HtrA) protein: Protease versus chaperone. IUBMB Life 68, 904–907 (2016).

61. S. Kim, R. A. Grant, R. T. Sauer, Covalent linkage of distinct substrate degrons controls assembly and disassembly of DegP proteolytic cages. Cell 145, 67–78 (2011).

62. H. Kim, K. Wu, C. Lee, Stress-Responsive Periplasmic Chaperones in Bacteria. Frontiers in Molecular Biosciences 8(2021).

63. W. M. Huston, C. Theodoropoulos, S. A. Mathews, P. Timms, Chlamydia trachomatis responds to heat shock, penicillin induced persistence, and IFN-gamma persistence by altering levels of the extracytoplasmic stress response protease HtrA. BMC Microbiol 8, 190 (2008).

64. G. Zhong, Chlamydia Trachomatis Secretion of Proteases for Manipulating Host Signaling Pathways. Frontiers in Microbiology 2(2011).

65. X. Wu, et al., The chlamydial periplasmic stress response serine protease cHtrA is secreted into host cell cytosol. BMC Microbiology 11, 87 (2011).

66. K. Hua, et al., Haemophilus parasuis Infection Disrupts Adherens Junctions and Initializes EMT Dependent on Canonical Wnt/β-Catenin Signaling Pathway. Front Cell Infect Microbiol 8, 324 (2018).

67. C. M. Overall, C. López-Otín, Strategies for MMP inhibition in cancer: innovations for the post-trial era. Nat Rev Cancer 2, 657–672 (2002).

68. S. Takeda, ADAM and ADAMTS Family Proteins and Snake Venom Metalloproteinases: A Structural Overview. Toxins (Basel) 8, 155 (2016).

69. Y. Okada, I. Nakanishi, Activation of matrix metalloproteinase 3 (stromelysin) and matrix metalloproteinase 2 (’gelatinase’) by human neutrophil elastase and cathepsin G. FEBS Lett 249, 353–356 (1989).

70. E. A. Garbett, M. W. Reed, N. J. Brown, Proteolysis in human breast and colorectal cancer. Br J Cancer 81, 287–293 (1999).

71. V. Noë, et al., Release of an invasion promoter E-cadherin fragment by matrilysin and stromelysin-1. J Cell Sci 114, 111–118 (2001).

72. Y. Ogata, J. J. Enghild, H. Nagase, Matrix metalloproteinase 3 (stromelysin) activates the precursor for the human matrix metalloproteinase 9. J Biol Chem 267, 3581–3584 (1992).

73. H. Nagase, R. Visse, G. Murphy, Structure and function of matrix metalloproteinases and TIMPs. Cardiovasc Res 69, 562–573 (2006).

74. W. C. Parks, C. L. Wilson, Y. S. López-Boado, Matrix metalloproteinases as modulators of inflammation and innate immunity. Nat Rev Immunol 4, 617–629 (2004).

75. W. Yan, S. X. Li, M. Wei, H. Gao, Identification of MMP9 as a novel key gene in mantle cell lymphoma based on bioinformatic analysis and design of cyclic peptides as MMP9 inhibitors based on molecular docking. Oncol Rep 40, 2515–2524 (2018).

76. H. G. Welgus, et al., Neutral metalloproteinases produced by human mononuclear phagocytes. Enzyme profile, regulation, and expression during cellular development. J Clin Invest 86, 1496–1502 (1990).

77. R. Kuefer, et al., ADAM15 disintegrin is associated with aggressive prostate and breast cancer disease. Neoplasia 8, 319–329 (2006).

78. A. J. Najy, K. C. Day, M. L. Day, ADAM15 supports prostate cancer metastasis by modulating tumor cell-endothelial cell interaction. Cancer Res 68, 1092–1099 (2008).

79. A. J. Najy, K. C. Day, M. L. Day, The ectodomain shedding of E-cadherin by ADAM15 supports ErbB receptor activation. J Biol Chem 283, 18393–18401 (2008).

80. R. de Waal Malefyt, J. Abrams, B. Bennett, C. G. Figdor, J. E. de Vries, Interleukin 10(IL-10) inhibits cytokine synthesis by human monocytes: an autoregulatory role of IL-10 produced by monocytes. J Exp Med 174, 1209–1220 (1991).

81. D. Y. Kim, K. K. Kim, Structure and function of HtrA family proteins, the key players in protein quality control. J Biochem Mol Biol 38, 266–274 (2005).

